# The dynamics of barrier loci recruitment along a continuum of divergence in the context of gene flow

**DOI:** 10.64898/2026.01.07.698084

**Authors:** Marjolaine Rousselle, Julie Jaquiéry, Jean Peccoud, Lucie Mieuzet, Frédérique Mahéo, Fabrice Legeai, Renaud Vitalis, Emmanuelle Jousselin, Jean-Christophe Simon, Mathieu Gautier, Carole M. Smadja

## Abstract

Understanding how reproductive isolation (RI) evolves in the context of gene flow is central to explaining how new species arise. Theory predicts that the accumulation of barrier loci depends on the interplay between divergent selection, recombination, and genomic architecture. However, empirical tests that track these processes across multiple stages of divergence within species remain scarce. The pea aphid complex, which comprises sympatric host-specialised biotypes spanning a continuum of divergence, provides a powerful opportunity to examine how ecological adaptation drives reproductive isolation despite ongoing gene flow. Using whole-genome sequencing data from 13 sympatric European biotypes, we constructed a dense, reference-anchored view of genomic divergence that controls for shared genetic background and recombination landscape. Joint analyses of genetic differentiation, absolute divergence, genetic diversity, recombination, and introgression rates reveal a consistent signature of divergence with gene flow and enable robust identification of barrier loci. Across the divergence continuum, RI is highly polygenic, but barrier loci are clustered and enriched in low-recombination regions and large chromosomal rearrangements, which are expected to strengthen linkage disequilibrium and promote the coupling of barrier effects. While genome-wide differentiation and the number of barrier loci increase gradually with divergence, differentiation within barrier loci displays patterns consistent with non-continuous dynamics, which could reflect threshold effects predicted by theory. Barrier loci contain excesses of salivary effector, detoxification and chemosensory genes, highlighting their central role in host plant specialisation and RI. The representation of these gene categories varies across divergence levels, and some loci are shared among independent biotype comparisons, suggesting common functional routes to specialisation via parallel evolution or introgression. Together, our results provide a comprehensive view of the genomic architecture and evolutionary dynamics underlying speciation with gene flow, illustrating how tightly linked, polygenic architectures can facilitate adaptation and diversification in natural populations.

## Introduction

Elucidating how species barriers emerge and evolve over time is crucial to explaining how new species arise, and therefore the origins of species diversity. Advances in speciation genomics have shown that speciation typically unfolds through the gradual accumulation of reproductive isolation (RI) loci across the genome and through time [1–3]. In the absence of complete geographic isolation between populations, this process can be counteracted by gene flow [4], leading to a loss of differentiation and the breakdown of reproductive barriers (e.g., [5]). However, another possible outcome in the context of gene flow is an increase in the strength of reproductive isolation and a progression towards more portions of the genome being protected from introgression [1]. Reinforcement is an example of one process that might be involved, where prezygotic isolation evolves in response to hybrid unfitness [4,6]. Because gene flow is common during population divergence [7], understanding how reproductive isolation evolves despite ongoing gene exchange remains a central challenge in speciation research [8–10].

Theory predicts that the outcome of divergence in the presence of gene flow depends mainly on the antagonism between selection and recombination among diverging loci, which influences how different reproductive barriers can evolve and get coupled to result in a stronger overall barrier to gene flow [4,11,12]. As a consequence, regimes of divergent selection, combined with mechanisms that locally reduce recombination, are key to fostering coupling between genetically independent barriers and the evolution of strong RI despite the disruptive effects of recombination [12,13]. Therefore, the maintenance and evolution of RI in the presence of gene flow is closely tied to its genetic architecture [14], and to mechanisms influencing recombination rate variation along the genome [15–17]. Chromosomal rearrangements, by reducing recombination in heterozygotes, have been proposed as one of the possible mechanisms promoting linkage disequilibrium and the concerted evolution of adaptive and barrier alleles [15,18].

Together, these considerations highlight the necessity of identifying the genomic architecture, functional nature, and timing of barrier loci to understand how reproductive isolation evolves and is maintained throughout the speciation process. Because speciation takes too long to observe directly, except in certain experimental evolution models [19], a widely adopted empirical approach to assess the dynamics of reproductive isolation is to examine the so-called “speciation continuum” or “divergence continuum” (i.e., sets of populations or species pairs showing varying degrees of RI) as a proxy for different stages of divergence [20]. This strategy has been applied to investigate the predictability of the speciation process, parallel evolution, and the build-up of barrier loci and genomic differentiation in various taxa including phytophagous insects forming host races [21–24], other insects (*Heliconius* butterflies [25,26], *Timema* stick-insects [27]), fish (threespine stickleback [28], cichlids [29]), birds (flycatchers [30]; silvereye [31]), marine snails (*Littorina saxatilis* [32,33]) and plants (monkeyflowers [34]; *Populus* trees [35]). While between-species comparisons within a clade reveal later stages of divergence, the early phases are better captured by population-level comparisons within a species (e.g., [31,33]). Most continuum studies aim to use independent lineage pairs, which help assess parallelism and shared genomic patterns across taxa (e.g., [29,32,36]). However, concerns have been raised regarding the use of this “continuum of divergence” framework to reconstruct the full “trajectory” of speciation, since it is unrealistic to assume that different pairs, representing different evolutionary contexts, followed and will continue to follow the same trajectory [3,20,25]. Here, we used as our prime comparative framework a refinement of the continuum approach, in which a single reference lineage served as the baseline for comparison with multiple sympatric lineages spanning a gradient of divergence, with all comparisons anchored to the same genetic background. This design controls for genomic background and recombination landscape, provides a consistent reference for mapping divergence, and allows assessment of the dynamics of RI along this contemporary gradient since genetic distance from the reference can be quantified [30,31,37]. It therefore enables a more direct comparison of the genomic landscape and functional nature of barriers to gene flow between the reference lineage and the others, at different levels of intraspecific divergence. We complemented this reference-based approach with the analysis of reference-free pairs of lineages enabling us to distinguish general speciation-related genomic patterns from signals driven by the evolutionary history of a single reference lineage.

As noted above, phytophagous insects forming partially reproductively isolated host races specialised on different plants have repeatedly provided the opportunity to examine the divergence continuum in the context of ecological speciation with gene flow [38]. The pea aphid species complex represents such a system. In Western Europe, 15 host races (or biotypes) have been identified, coexisting in sympatry and specialising on distinct Fabaceae plants [22,39,40]. Several lines of evidence indicate the evolution of some level of reproductive isolation among pea aphid biotypes driven by host-plant-induced divergent ecological selection. One component of reproductive isolation induced by host plant adaptation is selection against immigrants, evidenced by a lower performance (survival, fecundity) of specialised lines on non-host plants [22,41–43]. Host plant adaptation also induces host-dependent selection against hybrids, since F1 hybrids show lower performance on both parental host-plants, compared to the performance of each parent on its host-plant [44,45], with no indication of host-independent postzygotic isolation in the parthenogenetic phase of the pea aphid life cycle [45]. In addition, empirical evidence shows strong host plant choice: the different biotypes display a genetically based preference for the host plant from which they were collected [39,46,47]. These plants not only provide food resources, but also habitats and mating sites. Habitat choice and selection against immigrants are therefore thought to be the main drivers of assortative mating, since pea aphids reproduce on their host plants and there is no evidence for other prezygotic barriers so far [48]. Altogether, these results indicate that host plant specialisation, through both host plant adaptation and host plant choice, is the key component of reduced gene flow in pea aphids [49].

The extent to which pea aphid divergence has been sympatric remains uncertain; however, the biotypes are now broadly sympatric across Eurasia, and the presence of hybrids implies that they experience some level of ongoing gene flow. Despite this, reproductive isolation is maintained: the number of observed hybrids between two given biotype pairs in the field negatively correlates with their level of genetic differentiation, which suggests that pea aphid biotypes show different levels of reproductive isolation with one another, representing a continuum of RI between pairs of biotypes [20,22]. Previous results based on a restricted set of markers indicate that this continuum of divergence in the pea aphid spans a gradient of genetic differentiation (percentage of standardised within-group component of molecular variance *F*_SC_) ranging from 30 to 90%, and a gradient of observed proportion of F1 hybrids per biotype pair ranging from 0 to 3% [22]. The pea aphid complex thus provides an ideal system for examining how divergent selection can result in the build-up or maintenance of reproductive isolation in the face of gene flow along a gradient of divergence. Importantly, although the reference-based approach described above is seldom applicable because it requires multiple lineages that interact in nature, the pea aphid complex represents a rare natural system in which this framework can be implemented, owing to its gradual, intraspecific continuum of divergence among sympatric biotypes.

The pea aphid system also now benefits from the development of high-quality genomic resources and first insights into the genetics of reproductive isolation. Building on early QTL work on host-associated traits [42,50,51], several studies have since examined genomic differentiation and gene expression among biotypes, using a restricted number of markers [52,53], a limited number of biotypes [54] or candidate gene approaches [55–59]. These approaches revealed divergent genomic regions and genes among biotypes potentially acting as barriers to gene flow – notably genes belonging to functional categories expected to play a key role in host plant specialisation: chemosensory, detoxification and salivary effector genes. While chemosensory proteins (including receptors and binding proteins) allow aphids to recognise host-plant volatile and nonvolatile compounds, salivary effector proteins and detoxification enzymes (including P450 cytochromes) can help aphids to feed on a plant by circumventing its immune defences, promoting phloem sap uptake, and detoxifying plant secondary metabolites [59–61]. However, inference about the genetic architecture of barrier regions along the divergence gradient was limited by the use of a fragmented reference genome assembly [62] and a restricted range of biotypes [54].

Here, we combined the latest high quality, chromosome-scale assembly [63], a curated annotation of three categories of genes potentially involved in host plant specialisation [55,56,58,59,64–66], and whole-genome resequencing data from 13 sympatric biotypes spanning the whole continuum of divergence, to investigate patterns of reproductive isolation across different levels of divergence. Specifically, we inferred the genetic relationships between European biotypes and used this topology to estimate the age of divergence events and build reference-based pairs of biotypes of increasing genomic differentiation. We used this comparative framework to characterise the scenario of divergence, relying on the patterns of the genome-wide correlations between measures of differentiation and between- and within-population diversity [35,67,68]. This strategy allowed us to use measures of *F*_ST_, *D*_XY_ and *f*_dM_ [35,67–70] to identify putative regions porous vs. non-porous to gene flow. We then evaluated how the genomic architecture of barrier loci varies across different levels of divergence, focusing in particular on their number, size and clustering patterns, as well as their association with regions of reduced recombination and candidate chromosomal rearrangements. Finally, we explored the functional composition of barrier loci and patterns of candidate gene recruitment across the divergence gradient, focusing on several candidate gene families likely involved in host plant adaptation and in host plant choice.

## Results

### 1. Characterisation of the continuum of divergence and timing of divergence in the pea aphid complex

We focused this study on a set of pea aphid biotypes that span the whole continuum of divergence and occur in sympatry [22,40]. These constitute 13 European pea aphid populations specialised on distinct Fabaceae host species, and hereafter named after their host plants (e.g., the *Pisum sativum* biotype) (see Material & Methods and **Fig 1A**), for which we gathered whole-genome Pool-Seq data. To assess the relative genetic similarity among biotypes within the complex, we built a dendrogram based on genome-wide *F*_ST_ estimates for all pairs of European biotypes. This confirmed results previously obtained using a more restricted set of markers [22] (**Fig 1A**), notably that the different biotypes diverged largely in a successive, stepwise manner, rather than through hierarchical splitting from a small number of ancestral lineages, thus forming a continuum of divergence.

**Figure 1:**
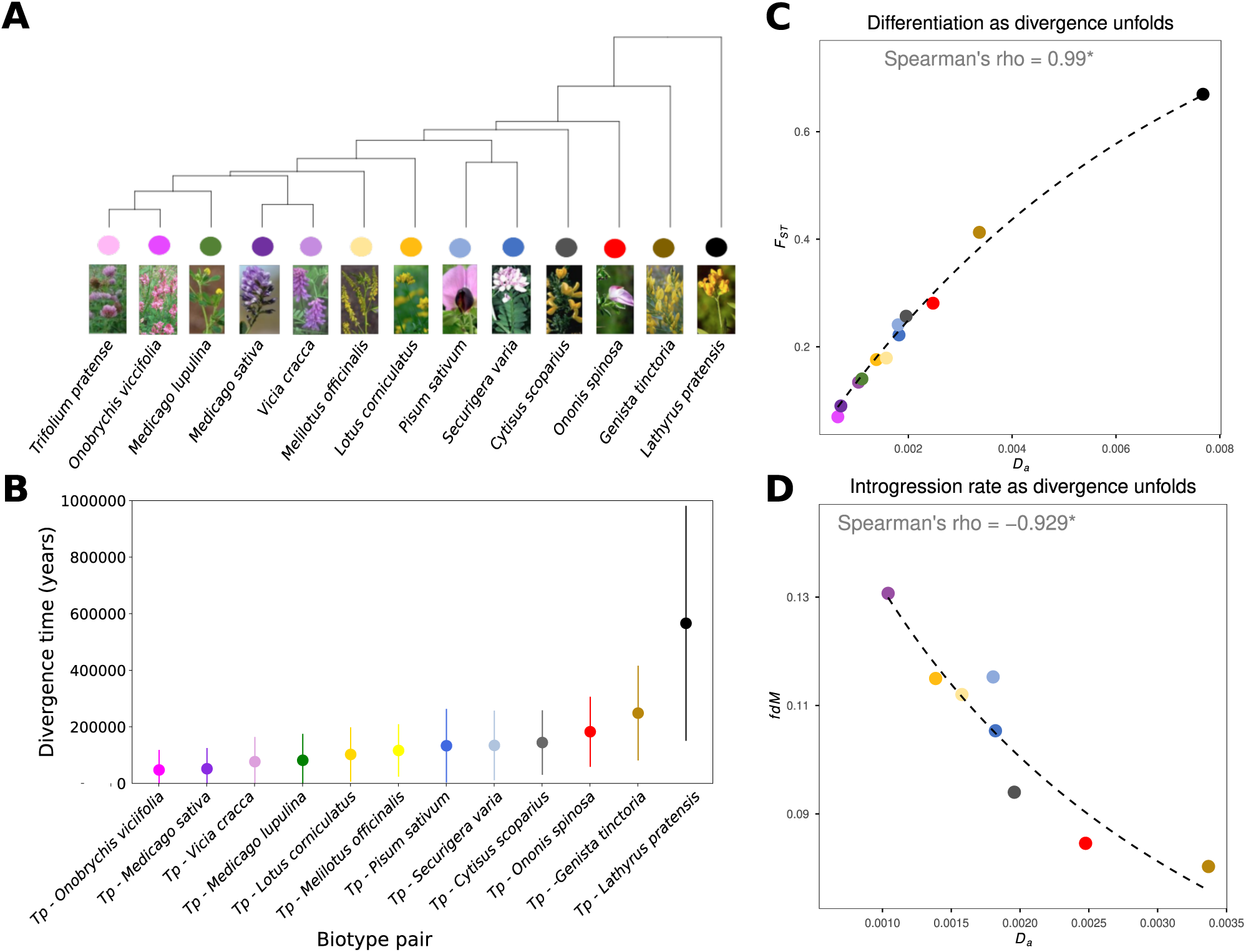
Characterisation of the continuum of divergence among European pea aphid biotypes. **A**: Topology of the relationships among 13 pea aphid biotypes, based on pairwise genome-wide *F*_ST_ estimates; **B**: Divergence times between the reference *Trifolium pratense* (*Tp*) biotype and the 12 other European biotypes obtained from rescaling net divergence (*D*_a_) values with estimate of the pea aphid mutation rate. Dots represent the different biotype pairs involving the *Trifolium pratense* biotype, with the different colours indicating the second biotype composing a pair, following the colour code used in panel A; **C**: Dynamics of genome-wide genetic differentiation (*F*_ST_) as *D*_a_ increases along the continuum of divergence. Colour codes follow panel B. The black dashed line represents the asymptotic regression fit (*b* = 143.5, *p* = 4.7 × 10^-14^); **D**: Dynamics of genome-wide introgression rate (*f*_dM_) (see Material and method for the computation of *f*_dM_ for some pairs of biotypes) as *D_a_* increases. Colour codes follow panel B. The black dashed line represents a hyperbolic Michaelis–Menten model fit (*b* = 448.5, *p* = 0.009), indicating rapid decay of introgression with increasing divergence.

To better understand the extent of gene exchange between these biotypes, we computed the *D*-statistic for all possible trios of biotypes, using as outgroup a biotype from Japan specialised on *Vicia cracca*, described as a putative cryptic species that does not exchange gene flow with the European lineages [71]. As a measure of precaution, we excluded from all analyses based on *D*-statistics, the biotype specialised on *Lathyrus pratensis* which may have diverged from the other European biotypes before this Japanese biotype (Jean Peccoud, personal communication). We then estimated *D*_min_, the minimum *D*-statistic for a given trio irrespective of the assumed underlying topology. We found that more than 96% of trios have significant *D*_min_, which implies reticulate evolution at multiple levels. To implement the framework laid out in the introduction, we established patterns of genomic differentiation and reproductive isolation between one reference biotype and the 12 others (Material and Methods). We chose the biotype specialised on *Trifolium* (clover) as reference, to maximise the range of divergence spanned by the 12 biotype pairs (**Fig 1A**).

Using a method similar to [72], based on estimates of net divergence (*D*_a_ – a measure of the divergence that accumulated since the split between the two biotypes) but revising the mutation rate estimate to account for both sexual and asexual reproduction during the aphid life-cycle (see [72] and Material & Methods), we first dated divergence times in the complex (**Fig 1B**). We estimate that the radiation dates back to a minimum of ∼545,000 years ago for the most distantly related biotype (*Lathyrus pratensis*), consistent with the results of [72]. Meanwhile, the closest biotype to *Trifolium pratense* (*Onobrychis viccifolia*) diverged at least 16,000 years ago.

Along the continuum of divergence, genome-wide *F*_ST_ correlates positively with *D*_a_ and increases asymptotically along the continuum of divergence (**Fig 1C**). For all correlations computed at the scale of the continuum, like this one, we performed equivalent tests without the *Lathyrus pratensis* biotype to ensure that the significance level and trends were not solely driven by this distantly related biotype. Unless specified otherwise, removing this biotype did not change the significance of correlations at the 5% risk (**S1 Table**).

We then computed the genome-wide average introgression rates between the reference *Trifolium pratense* biotype and the other biotypes by calculating the *f*_dM_ statistic along the genome [70] in trios that include biotype pairs of the continuum, where this measure is informative of potential gene flow (see Material and methods). This was feasible for eight out of 12 biotype pairs (see Material and methods). We found that the introgression statistic *f*_dM_ decreased monotonically with increasing net divergence (*D*_a_) and a hyperbolic model fit the data better than an asymptotic model or an exponential decay model (AIC_hyperbolic_ = - 55.5; AIC_exponential_ = −54.1; AIC_asymptotic_ = −20.7, **Fig 1D**), capturing the progressive reduction in gene flow as divergence accumulated. This further confirms that European biotypes are connected by gene flow, the amount of which decreases over time as reproductive isolation becomes stronger.

### 2. Genomic differentiation and diversity landscapes consistent with a scenario of divergence with gene flow

Understanding the genetic basis of reproductive isolation in pea aphid requires identifying genomic regions that are resilient to gene flow. Recent studies have made clear that the so-called genomic “islands of differentiation”, i.e., genomic windows with high levels of genetic differentiation (*F*_ST_) relative to the rest of the genome, do not necessarily represent loci involved in reproductive isolation or ecological specialisation, but can arise under several alternative selection-based scenarios [67,68,73]. To disentangle such scenarios and assess if the genetic differentiation, divergence and diversity landscapes are consistent with a scenario of divergence with gene flow in the pea aphid complex, we first computed genome-wide correlations between *F*_ST_, *D*_XY_ and genetic diversity, *π*, over 50kb non-overlapping windows in all pairs along the continuum (**S2 Table**, **Fig 2A**) and compared the observed correlation patterns with those expected under different scenarios of divergence [67,68,73].

**Figure 2:**
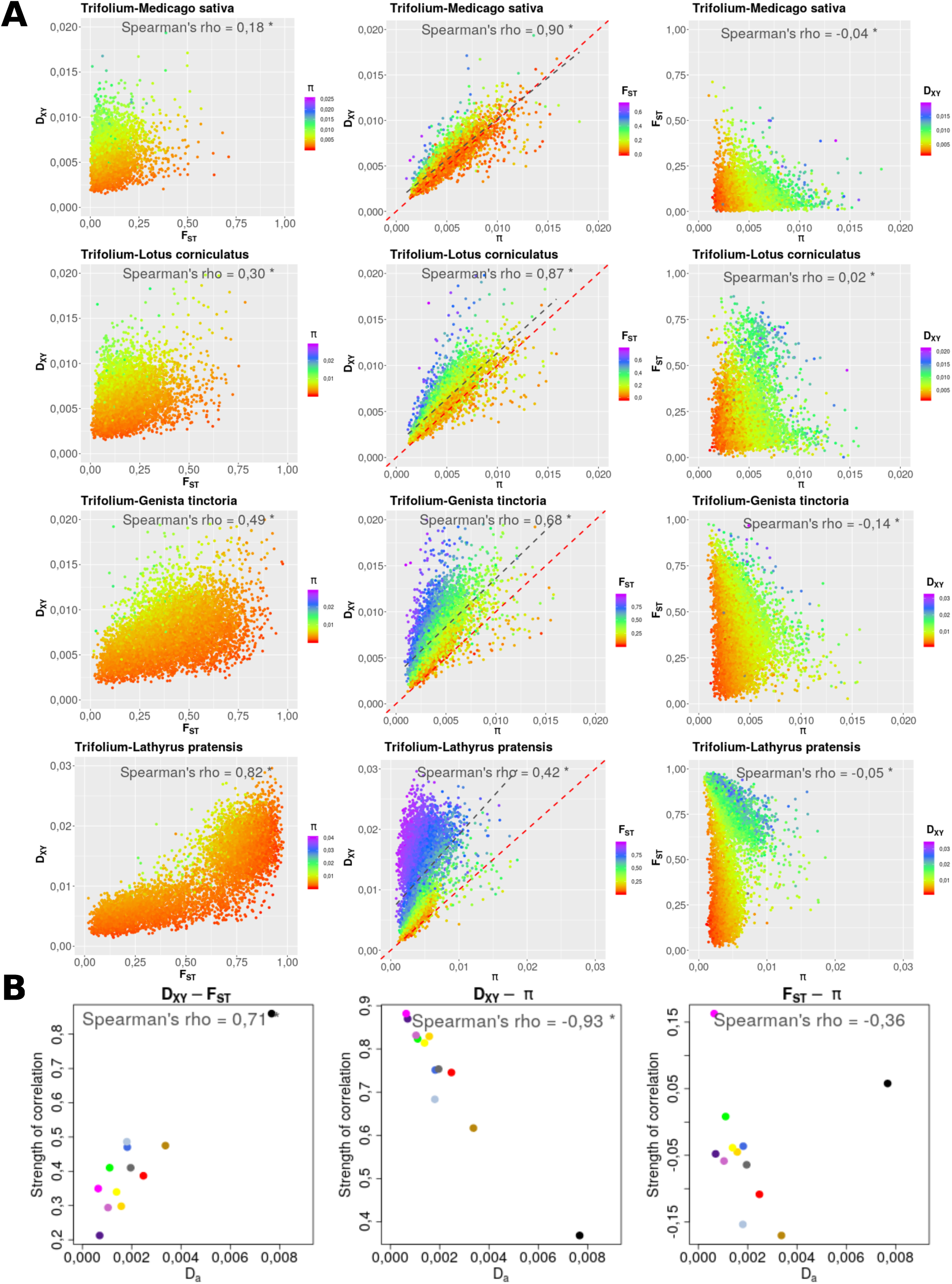
Genome-wide correlations between divergence and diversity statistics along the continuum of divergence. **A**: Correlations between *F*_ST_, *D*_XY_ and *π* in four biotype pairs of increasing divergence along the continuum. In each plot, dots represent the different genomic windows for which the different statistics could be computed. Grey dotted lines represent linear regression fits and red dotted lines represent identity lines. **B**: Dynamics of the strengths of correlations between diversity and divergence statistics as net divergence increases along the continuum. Dots represent the 12 different biotype pairs constructed using the *Trifolium pratense* biotype as reference, with the different colours indicating the second biotype composing a pair, following the colour code used in Fig 1.

We found that *F*_ST_ and *D*_XY_ estimates are significantly positively correlated in all biotype pairs (Spearman correlation coefficient ranging from 0.18 to 0.82) (**Fig 2A, S1 Fig**), and those correlations strengthen as net pairwise divergence increases (**Fig 2B**). This is not consistent with a simple scenario of recurrent positive or background selection, but is expected if the heterogeneous *F*_ST_ landscape is primarily explained by some regions being resistant to gene flow whereas the genomic background is not. *D*_XY_ and *π* are also strongly positively correlated, and as pairwise divergence increases, *D*_XY_ becomes increasingly higher than *π* (**Fig 2A, S2 Fig**) and the correlation between the two weakens dramatically (**Fig 2B**). This is consistent with a scenario where *D*_XY_ is initially affected by the ancestral population diversity, but as population divergence and reproductive isolation increases, ancestral diversity impacts *D*_XY_ less than does the accumulation of differences in regions resistant to gene flow. We observed that *D*_XY_ becomes comparatively higher than *π* especially in windows of high *F*_ST_ (**Fig 2A**). All these correlations hold when excluding the most divergent biotype, specialised on *Lathyrus pratensis*, from the analyses (**S1 Table**). In addition, all those patterns also hold when analysing a set of reference-free (i.e., without the *Trifolium pratense* biotype in common) biotype pairs of increasing divergence (*Trifolium pratense* vs*. Onobrychis viciifolia, Medicago lupulina* vs*. Vicia cracca, Pisum sativum* vs*. Melilotus officinalis, Ononis spinosa* vs*. Genista tinctoria*) (**S3** and **S4 Figs**). Thus, correlations, and their increasing strength as net divergence increases, are not driven by the non-independence of reference-based biotype pairs nor are they due to the use of the *Trifolium* biotype as a reference. Together, the correlation patterns between *F*_ST_ and *D*_XY_ and between *D*_XY_ and *π* are consistent with a scenario of divergence with gene flow where loci contributing to reproductive isolation restrict gene exchange between diverging species [67,68,73].

*F*_ST_ and *π* are mostly weakly negatively correlated (**Fig 2A, S5 Fig**), and those correlations do not significantly strengthen or weaken as divergence increases (**Fig 2B**). However, when excluding the *Lathyrus pratensis* biotype, this correlation becomes significantly negative (**S1 Table**). This pattern is not associated with low *D*_XY_ in high *F*_ST_ regions, which rules out a predominant scenario of allopatric recurrent selection, as we would expect high *F*_ST_ in low *π* regions, as observed, but not higher *D*_XY_. To further assess the potential influence of linked selection on patterns of diversity and differentiation, we evaluated the relationships between those statistics, and statistics that correlate with linked selection: recombination rate and gene density, (see Material & Methods). We found that gene density is negatively correlated to *π* (Spearman correlation coefficient ranging from −0.02 to −0.26, **S6 Fig**) and *D*_XY_ (Spearman correlation coefficient ranging from −0.05 to −0.16, **S7 Fig**), except for *Lathyrus pratensis.* This is consistent with the recurrent action of linked selection. However, under this model, *F*_ST_ is expected to show a positive relationship with gene density, whereas we found only very weak, often negative correlations (Spearman correlation coefficient ranging from −0.06 to 0.08, **S8 Fig**). Recombination rates yielded weak positive correlations with genetic diversity (**S9 Fig** and *D_XY_***S10 Fig**). *F_ST_* and recombination rate are mostly weakly negatively correlated (**S11 Fig**), and those correlations did not become stronger with divergence, which is inconsistent with expectations under the recurrent action of background selection [36]. Those different results illustrate the weak influence of recurrent linked-selection on the heterogeneous patterns of diversity and divergence in the pea aphid complex.

Finally, in the eight biotype pairs where we could estimate *f*_dM_, we found significantly negative correlations between *f*_dM_ and *F*_ST_ (Spearman correlation coefficient ranging from - 0.10 to −0.71) and between *f*_dM_ and *D*_XY_ (Spearman correlation coefficient ranging from - 0.05 to −0.64) (**S12** and **S13 Figs**). Both relationships became stronger as pairwise divergence increased (**S14 Fig**). This constitutes additional support for a scenario of divergence with gene flow, where regions of elevated *F*_ST_ and *D*_XY_ are primarily driven by reproductive isolation while recent or current gene flow affects the rest of the genome.

### 3. Genomic architecture of barrier loci along the continuum and the role of inversions

Given the results described above, we could identify barrier loci as 50-kb genomic windows showing both significantly elevated *F*_ST_ and *D*_XY_ compared to the rest of the genome [35,67,68] (see Material & Methods). Combining those two statistics is a conservative approach to identify regions of reduced gene flow causing both higher *F*_ST_ and higher average coalescent time, and not simply regions of elevated *F*_ST_ that could arise as a result of recurrent selection [68]. After merging adjacent genomic windows identified as barrier loci (called “barrier regions”), we found between 175 and 388 barrier regions across the different biotype pairs, representing between 3.5% and 14% of the genome (depending on the pair) and distributed across all four chromosomes (**S2 Table** and **Fig 3**). As expected, barrier loci showed lower introgression rate (*f*_dM_) than non-barrier loci, this pattern becoming stronger as divergence increased (**S15 Fig**). This emphasises that the combination of elevated *F*_ST_ and *D*_XY_ to identify barrier loci is reliable, even if a few regions defined as barrier loci show a rather elevated *f*_dM_, which could be attributed either to noise in the estimation of *f*_dM_ or to mis-identification of barrier loci. Overall, we characterise a highly polygenic architecture of reproductive isolation in the pea aphid.

**Figure 3:**
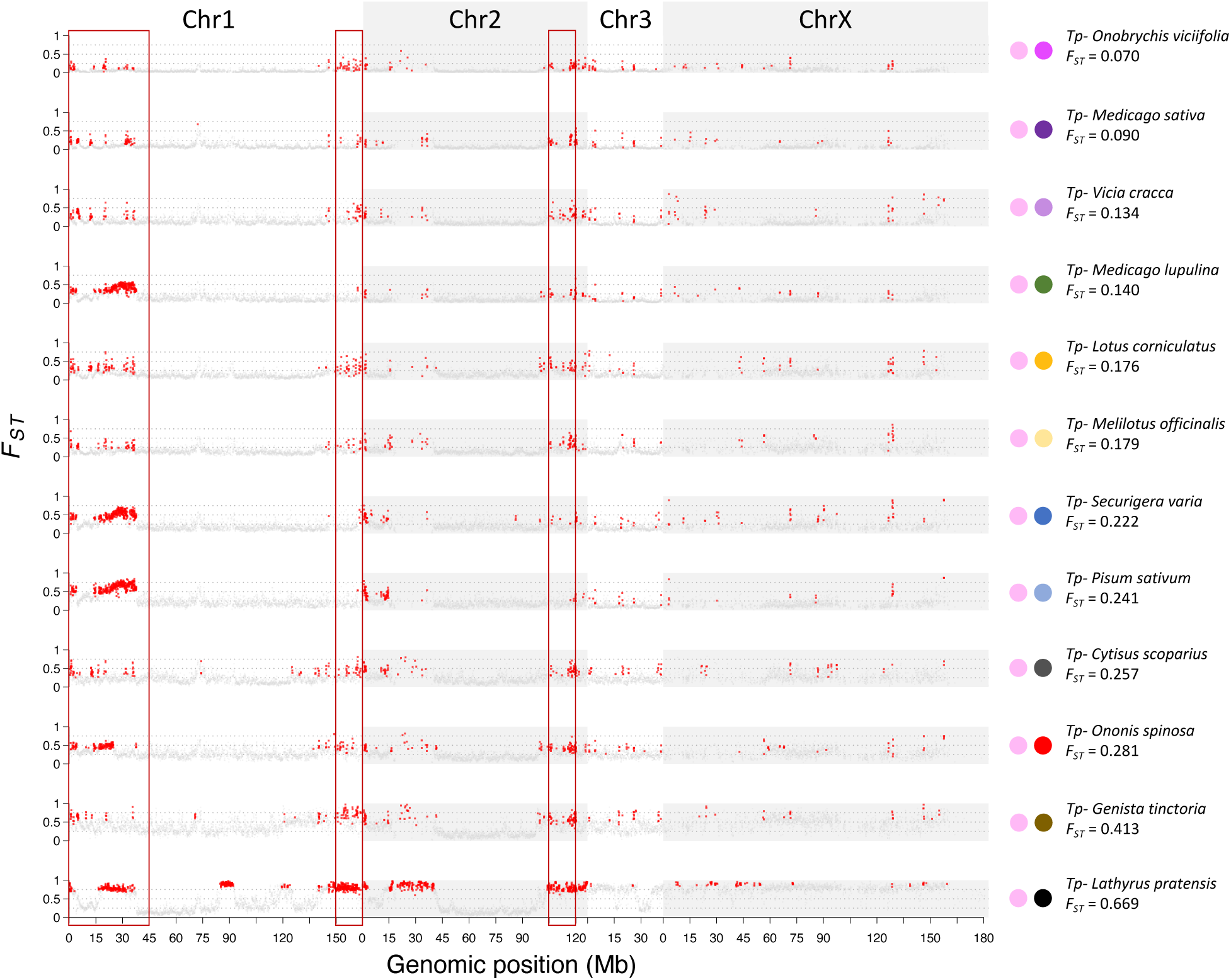
Genomic landscapes of *F*_ST_ and barrier loci between the *Trifolium pratense (Tp)* biotype and the other biotypes along the continuum of divergence. Red dots represent barrier loci, i.e. 50 kb genomic windows showing both significantly elevated *F*_ST_ and *D*_XY_; grey dots represent non-barrier loci. Red rectangles represent genomic regions with a significant enrichment of barrier loci detected in multiple biotype pairs (here in at least seven biotype pairs). Biotype pairs are ordered by increasing genome-wide *F*_ST_.

The total length of barrier loci (**Fig 4A**) and the number of (merged) barrier regions (**Fig 4B**) significantly increases with net divergence *D*_a_ across the continuum of divergence. However, we did not find a significant correlation between the average length of barrier regions and net divergence (**Fig 4C**). Surprisingly, we detected that three biotype pairs with intermediate levels of genetic differentiation display particularly elevated average lengths of barrier loci (*Pisum sativum*, *Securigera varia*, and *Medicago lupulina*). Finally, interesting relationships between genetic differentiation (*F*_ST_) and net divergence at barrier loci and non-barrier loci emerged from non-parametric cubic regression spline (CRS) models. For barrier loci, CRS selected a piecewise-linear model with five segments (degree = 1), corresponding to four visually distinct divergence regimes (**Fig 4D**, circular points). This model fitted the data extremely well (*R^2^* = 0.990, residual SE = 0.00025, cross-validation score = 1.92×10⁻⁷), and was strongly favoured over the asymptotic model (AIC_spline_ = - 207.08 vs AIC_asymptotic_ = −64.79). By contrast, non-barrier loci were best described by a single polynomial segment model (degree = 4, 1 segment) with a lower AIC than the corresponding asymptotic model (AIC_spline_ = −229.21 vs AIC_asymptotic_ = −81.99), indicating a smooth and continuous increase (**Fig 4D**, diamond points; *R^2^* = 0.9985, residual SE = 0.00093, CV score = 5.53 x 10⁻⁷), similar to genome-wide *F*_ST_ patterns (**Fig 1C)**. Thus, barrier loci exhibit structured, threshold-like changes in genetic differentiation as divergence from the *Trifolium* biotype increases, whereas differentiation at non-barrier loci changes gradually. Although the analyses were based on a limited number of biotype pairs (*n* = 12) and the number/position of spline segments should be interpreted cautiously, the low cross-validation scores and lack of overfitting indicate that the qualitative contrast between barrier and non-barrier divergence patterns is robust.

**Figure 4:**
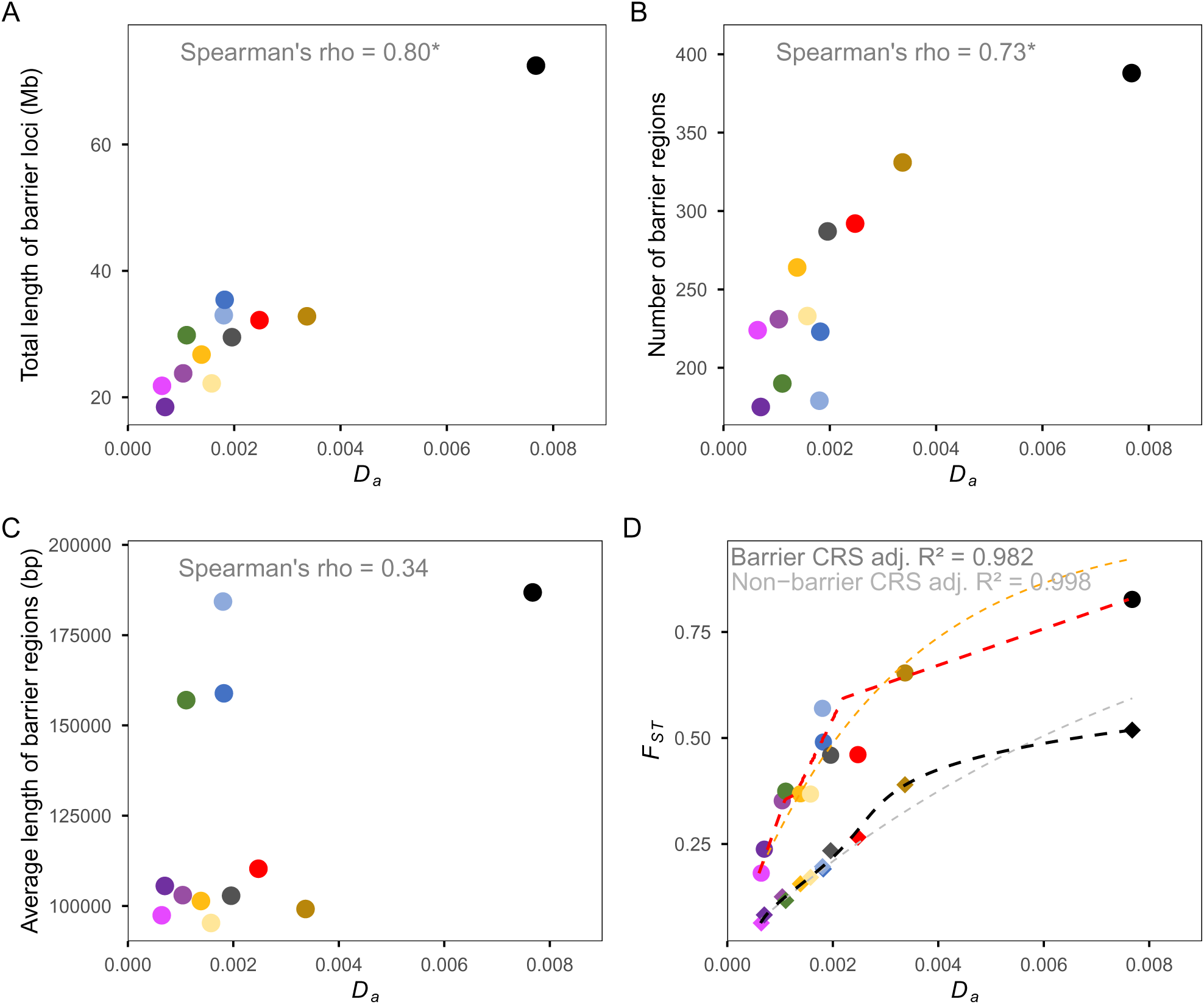
Characteristics of genomic regions acting as barriers to gene flow as divergence increases across the continuum. Coloured dots represent biotype pairs as in Fig 1B. **A**: Total length of barrier regions (i.e., merged adjacent barrier loci) as net divergence ( *D*_a_) increases (in Mb), **B**: Number of (merged) barrier regions as net divergence increases, **C**: Average length of (merged) barrier regions as net divergence increases, **D**: Mean *F_ST_* within barrier windows as net divergence increase (circular points) and mean *F*_ST_ in non-barrier windows (randomly chosen among all non-barrier windows to obtain the same number of non-barrier windows as the number of identified barrier windows, for each biotype pair) as divergence increases (diamond points). Red (for barrier loci) and black (for non-barrier loci) dashed lines show regression spline (CRS) fits; orange (for barrier loci) and grey (for non-barrier loci) dashed lines show asymptotic fits.

While barrier loci are widely distributed along the genome, each biotype pair presents genomic regions with significant enrichment of barrier loci compared to a null distribution (permutation tests, **S3 Table**). Notably, several of these regions are shared by most biotype pairs along the continuum (**S3 Table, Fig 3**). This suggests that these genomic regions repeatedly accumulated barrier loci isolating the *Trifolium* biotype and the others, regardless of their level of divergence. This trend also applies to our set of four independent pairs of biotypes (*Trifolium pratense* vs*. Onobrychis viciifolia, Medicago lupulina* vs*. Vicia cracca, Pisum sativum* vs*. Melilotus officinalis, Ononis spinosa* vs*. Genista tinctoria*) as they display very similar landscapes of barrier loci along the genome (**S16 Fig)**. This result indicates that the observed patterns of barrier loci distribution in the reference-based biotype comparisons do not strongly depend on the choice of the *Trifolium* biotype as a reference. Among all barrier loci characterized in the four independent biotype pairs, we identified 24 barrier loci shared by all four pairs, on average 50 shared by three pairs, on average 35 shared by two pairs vs. on average 104 that are specific to a biotype pair (**S17 Fig**).

Across all biotype pairs examined, between 12 and 43% of barrier loci were concentrated in the first ∼40Mb of chromosome 1 (**Fig 3** and **S16 Fig**). Alignment of two genome assemblies (one from a *Medicago sativa* individual (AL4f) [74], and the reference genome we used in this study, from a putative *Pisum sativum* individual (JIC1)) indicates that this region corresponds to large structural differences, consistent with one translocated and two inverted segments (**Fig 5A**). Using a method recently proposed for Pool-Seq data [75], we inferred that this rearranged region segregates at different frequencies in most biotypes of the continuum (see Material and methods). Interestingly, the proportion of barrier loci located within this rearranged region among all barrier loci in a given biotype pair significantly negatively correlates with the frequency of the rearrangement in the non-*Trifolium* biotype forming the pair (**Fig 5B**).

**Figure 5:**
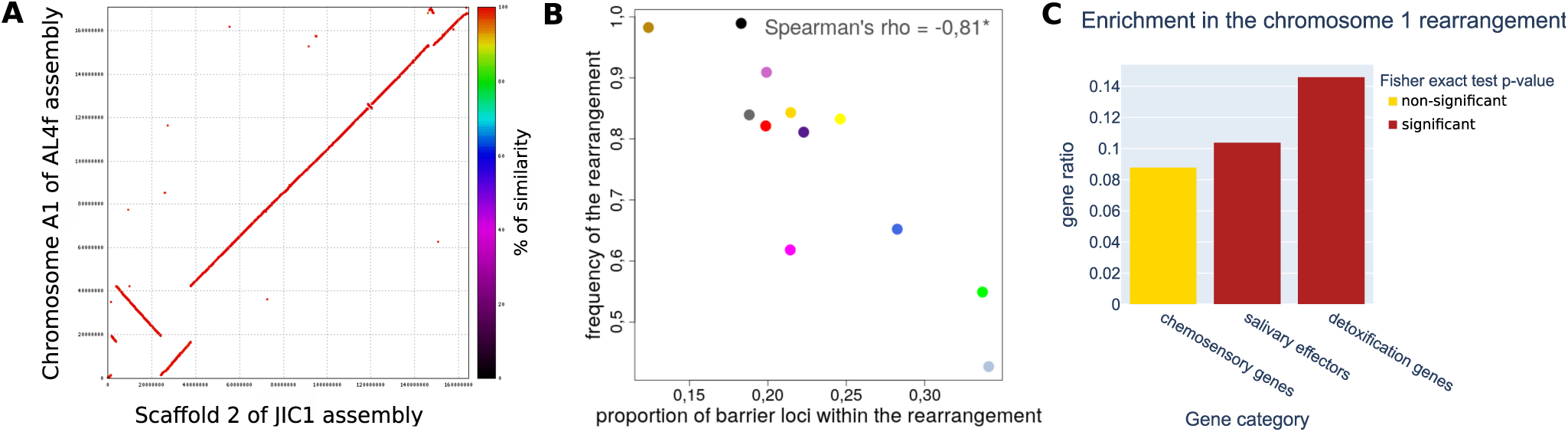
Analysis of barrier loci and gene content within a large chromosomal rearrangement on chromosome 1. **A**: Dot plot of chromosome A1 from an individual specialised on *Medicago sativa* AL4f (GCA_005508785.2) (Li et al. 2019) vs. scaffold 2 of the genome assembly of an individual putatively specialised on *Pisum sativum*, JIC1 (Mathers et al. 2021); **B**: Relationship between the frequency of the rearrangement in different biotypes and the proportion of barrier loci within this region in the different biotype pairs – coloured dots represent biotype pairs as in Fig 1; **C**: Fisher exact test results denoting the overrepresentation of genes annotated as chemosensory genes, salivary effectors and detoxification genes within the rearranged region. Gene ratio represents the ratio of the number of outlier genes (i.e., genes within barrier loci) of each category within the inversion over the number of outlier genes of each category outside the inversion.

Strikingly, the three biotypes where the rearrangement segregates closest to intermediate frequency (*Pisum sativum*, *Securigera varia*, *Melilotus officinalis*) also harbor distinctly large barrier loci (**Fig 4C**), a signal which disappears when we exclude this rearranged region (**S18 Fig**). This indicates that barrier loci located within this rearrangement are particularly large when it segregates at intermediate frequencies (**S19 Fig**). With respect to gene families hypothesised to play a role in host plant specialisation (see Introduction and next section), we found that this region harbors more genes identified as salivary effectors in previous studies on the pea aphid [76,77] or annotated as detoxification genes (e.g., P450, GSTs) than expected by chance (**Fig 5C**).

Beyond this large rearranged region on chromosome 1, we identified other genomic rearrangements based on the alignment of the two versions of the pea aphid genome: we found an additional large (∼17Mb) translocated region at the distal end of chromosome 1, another large rearrangement made of two consecutive inversions located in the first 15Mb of chromosome 2, a roughly 2Mb inversion on chromosome X and seven other smaller rearrangements/inversions (<2Mb) on chromosomes 1, 2 and 3 (**Fig 5A**, **S20 Fig**). We found that barrier loci overlap with those rearrangements significantly more than expected by chance (permutation tests p-values are all below 0.005 for all pairs but the *Genista tinctoria* vs*. Trifolium pratense* pair (*p*-value = 0.069)). At the scale of the whole genome, we found that barrier loci were on average more associated with lower recombination rate than non-barrier loci, with the exception of the two most divergent pairs involving *Genista tinctoria* and *Lathyrus pratensis* biotypes (**S21 Fig**). If our identification of barrier loci were induced by background selection rather than reproductive isolation, this pattern should be exacerbated in windows of high gene density where stronger linked purifying selection would influence *F*_ST_. However, the association between barrier loci and low recombination rates appears less significant in windows of high gene density compared to those of low gene density (**S21 Fig**), supporting the robustness of our inferences. Additionally, this result does not seem to be due to a detection bias, as we did not observe higher variance in *F*_ST_ and *D*_XY_ among genomic windows in low recombination regions (**S22 Fig**).

### 4. Dynamic recruitment of the molecular actors involved in host plant specialisation along the continuum

We next analysed the gene content of inferred barrier loci. Across all biotype pairs, we identified between 1,136 (in the *Trifolium pratense* vs*. Medicago sativa* pair) and 4,479 genes (in the *Trifolium pratense* vs*. Lathyrus pratensis* pair) within barrier loci, with a mean+/-SE of 1,984 +/- 204, out of more than 30,000 annotated genes. We hereafter refer to these as “outlier” genes (the complete list of which is reported in **S4 Table**). A substantial proportion of these genes (65% on average) are located in genomic windows identified as highly or very highly significant barriers in at least one biotype pair (i.e., barriers with both significantly elevated *D*_XY_ and *F*_ST_, at the alpha-level 0.01 or 0.001 of the permutation tests) (**S4 Table**). Although some outlier genes may show elevated divergence between biotypes due to hitchhiking effects without directly impacting gene flow, the most divergent ones may present variants that play a direct role in reducing gene flow and/or are under divergent selection.

We focused our functional analyses on three gene categories with available high-quality annotation (see Material and Method) and which are known or expected to play roles in host-plant specialisation in pea aphids or other phytophagous insects [60,61,78]. One gene category comprises chemosensory genes, potentially involved in host plant choice, with some genes already identified as outliers in previous candidate gene approaches and QTL analyses [55,56,58,79]. They include genes coding for Olfactory Receptors, Gustatory Receptors, Ionotropic Receptors, Odorant Binding Proteins, ChemoSensory Proteins, and Sensory Neuron Membrane Proteins [64–66]. Another category comprises detoxification genes, potentially involved in dealing with harmful plant metabolites, including cytochrome P450 genes, glutathione S-transferase (GST) genes and UDP-Glycosyltransferase (UGT) genes [61,80]. A third category represents genes identified as potential salivary effectors in the pea aphid through experimental work and involved in responses to plant defense [59,76,77,81].

All three gene categories proved highly significantly overrepresented among outlier genes (*n* = 37 outlier detoxification genes, Fisher exact test *p*-value = 0.0022;; *n* = 71 outlier chemosensory genes, Fisher exact test *p*-value = 0.0026; *n* = 1,432 outliers among potential salivary effector genes, Fisher exact test *p*-value < 2.2e-16, **S4 Table**). This significant overrepresentation was found in most of the biotype pairs involving the *Trifolium* biotype, with some exceptions (**S23 Fig**). Excess of chemosensory genes was highly significant, except for the *Medicago lupulina*, *Pisum sativum* and *Securigera varia* pairs, while detoxification genes were overrepresented among outliers only in the *Medicago lupulina*, *Pisum sativum, Ononis spinosa*, *Genista tinctoria* and *Lathyrus pratensis* pairs.

Because extended linkage and hitchhiking, particularly among genes belonging to multigene families, may inflate the number of outlier genes, we sought to refine the list of candidate genes by focusing on those most likely to present variants underlying reproductive isolation. Specifically, we identified outlier genes located within the most differentiated and divergent barrier loci (“highly” or “very highly significant” loci)) and consistently detected across a majority of biotype pairs along the continuum of divergence (see Material and Method). These 556 top outlier genes, representing ∼6% of all outlier genes (**S4 Table**), included genes lying within the large rearranged region on chromosome 1 (30% of the top outliers), among which 28 were potential salivary effectors and four were chemosensory genes – OBP13, Or21, Or22 and Or3. Among all top outlier genes, we identified 90 putative salivary effector genes, including one sucrase gene (S1) and one GST detoxification gene. Some chemosensory genes were also among those showing the strongest and most consistent between-biotype differentiation along the continuum of divergence. Notably, they span all known chemosensory gene families: seven Gustatory Receptor genes (Gr7, Gr8, Gr9, Gr10, Gr38, Gr39 and Gr40), six Odorant Receptor genes (Or3, Or11, Or12, Or21, Or22 and Or28), one Ionotropic Receptor gene (Ir327), one Odorant Binding Protein (OBP13) and one Chemosensory Protein (CSP10). Interestingly, top outlier genes were strongly clustered along the genome, with a median nearest-neighbour distance among top outliers of 7.7 kb (± 22.5 kb) and 60.8% of genes having a nearest-neighbour distance ≤ 10 kb, compared with 2.8% expected by permutation (*p* <0.001, **S24A-B Fig**). This significant physical clustering was also found outside the candidate chromosomal rearrangements (57% of genes with a neareast-neighbour distance ≤ 10 kb, permutation test: *p* < 0.001, **S24C-D Fig**), supporting a biological signal beyond structural context. This tight physical linkage involves genes belonging to different gene families (e.g., Or and OBP outlier genes) or different categories of candidate genes (e.g., chemosensory and salivary effector outlier genes, see **S4 Table** and **Fig 6**). To evaluate whether any of these prime candidate genes might be repeatedly involved in the divergence of several biotype pairs – potentially representing shared reproductive isolation genes and/or common functional determinants of host plant specialisation – we searched for outlier genes detected in all four independent biotype pairs and showing the strongest signals of divergence in these pairs. Only a small subset (66, ∼12%) of the prime candidates met these criteria (**S4 Table**), including 15 potential salivary effector genes and two chemosensory genes, Odorant Receptors Or3 and Or28.

**Figure 6:**
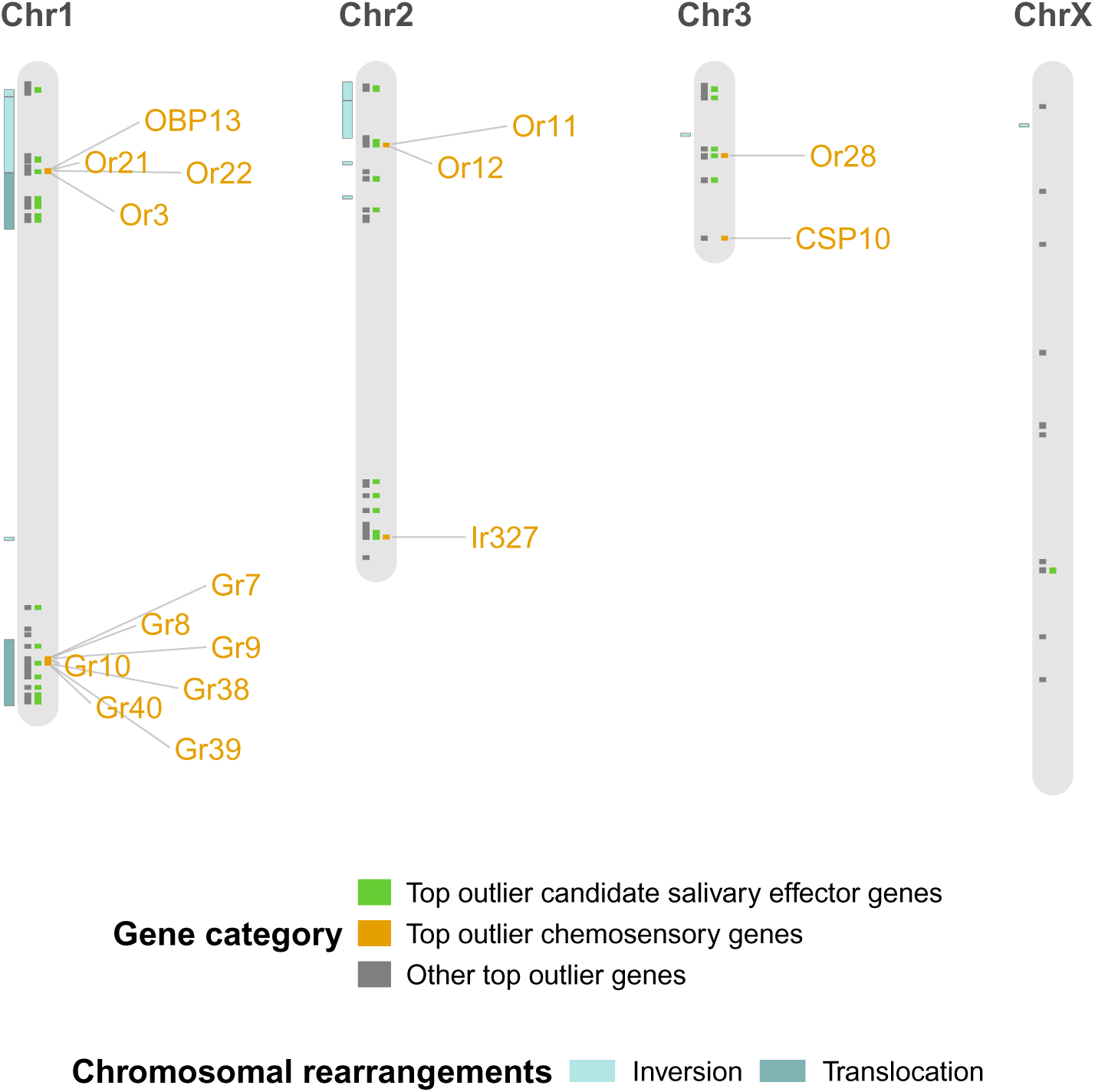
Genomic distribution and nature of top outlier genes identified across the continuum of divergence. Top outlier genes are represented as small rectangles according to their position within each chromosome, and the gene category they belong to (candidate salivary effector genes, chemosensory genes or other genes) represented in different colours and horizontal tracks. Genomic positions of identified candidate chromosomal rearrangements and the names of top chemosensory genes are specified on the left- and right-hand sides of each chromosome.

To assess the dynamics of outlier gene recruitment during biotype divergence, we characterised the variation in the number of outlier genes as a function of net divergence. We found that the number of outlier genes within salivary effector and detoxification candidate gene categories significantly increased with divergence from the reference biotype on *Trifolium pratense* (**Fig 7D, G**). We also found a trend towards a lower proportion of chemosensory and salivary effector outlier genes as divergence increases (**Fig 7B, E**), which could suggest a dilution effect if genes from other functional categories contribute to differentiation in more divergent biotype pairs. These other gene categories may either underlie other reproductive barriers arising later in the speciation process, or coupling processes may extend divergence to neutral regions of the genome. No gene category consistently showed higher between-biotype differentiation than the others when it came to the most closely related biotypes, but some showed a high number of outlier genes in some of these biotype pairs (e.g., chemosensory genes in the *Onobrychis viciifolia* biotype pair, salivary effectors and detoxification genes in the *Medicago lupulina* biotype pair). We thus classified outlier genes according to their occurrence among barrier loci identified in the less divergent biotype pairs or in the more divergent ones, and we used this information as a proxy to their contribution to early or late divergence (see Material and Methods). We found that chemosensory genes were significantly overrepresented in both “early” and “late” barrier loci, while salivary genes were only overrepresented in “early” barrier loci and detoxification genes were only overrepresented in “late” barrier loci (**Fig 7C, F, I**).

**Figure 7:**
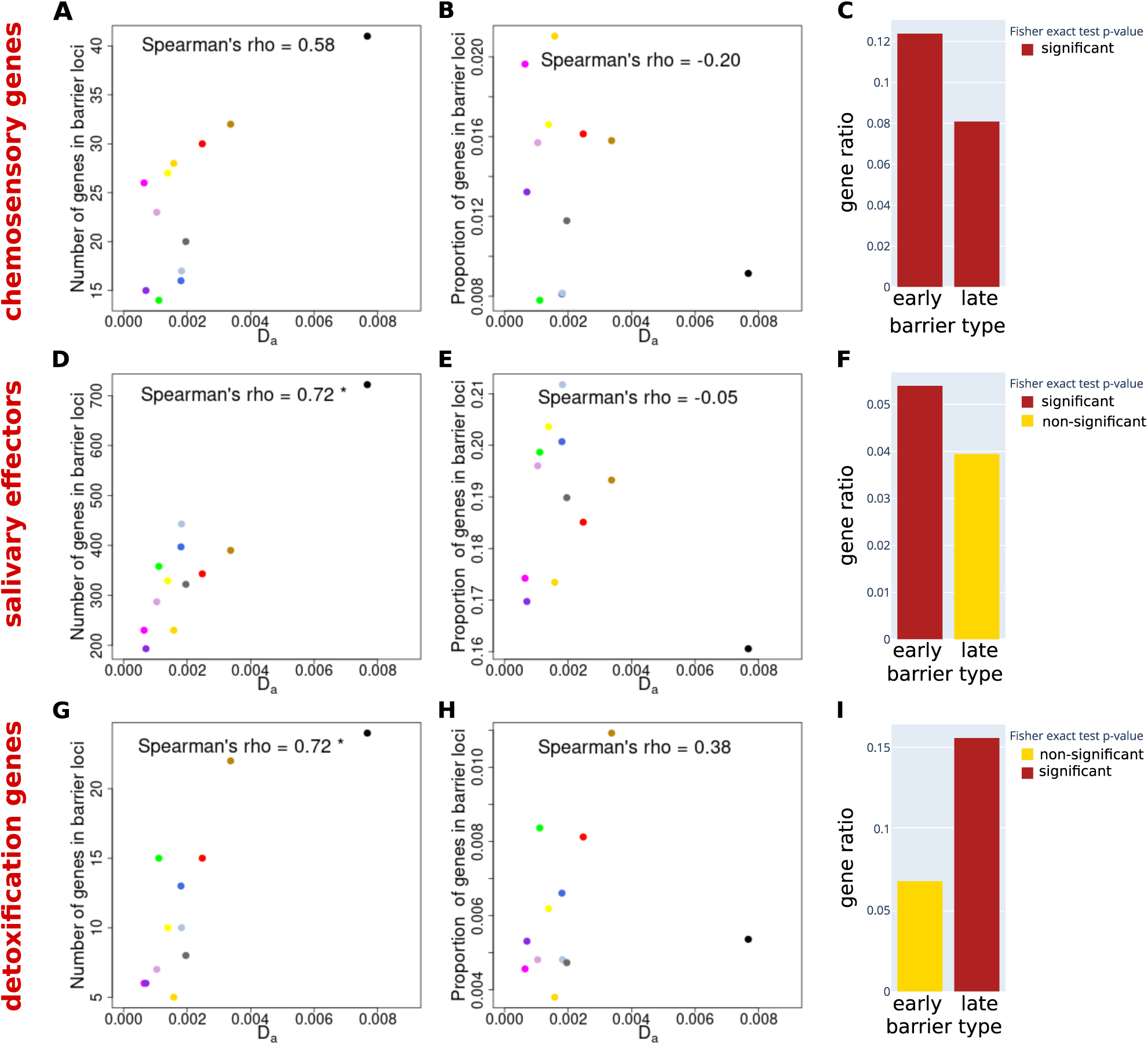
Dynamics of outlier genes belonging to the chemosensory, potential salivary effector and detoxification gene categories as divergence increases along the continuum of divergence. **A,D,G**: Number of the three categories of candidate genes within barrier loci across the continuum. **B, E, H**: Proportion of genes belonging to the three categories of candidate genes within barrier loci compared to the total number of genes within barrier loci across the continuum. **C, F, I**: Overrepresentation of the three categories of candidate genes among “early” or “late” barrier loci. Gene ratio represents the ratio of the number of genes of each category within early or late barrier loci over the number of genes of this category outside of early or late barrier loci.

## Discussion

The pea aphid is a classic model of ecological speciation, characterised by multiple host-plant specialised biotypes in which adaptation to distinct legumes, coupled with mating on these plants, generates reproductive isolation [49]. This system provides a unique opportunity to study reproductive isolation across a continuum of divergence, from closely-related biotypes with extensive ongoing gene flow to lineages that are almost fully isolated [22,49]. Using genome-wide data from 13 sympatric European biotypes, we characterised patterns of differentiation and reproductive isolation across this continuum. Confronting all biotypes to a reference biotype brought insights into the nature and genomic architecture of barriers to gene flow across different stages of divergence. The genetic architecture of reproductive isolation identified by this reference-based approach are not specific to the evolutionary history of the *Trifolium*-specialised biotype, as several barrier loci appeared shared by four independent biotype pairs, consistent with parallel recruitment and/or introgression of alleles underlying ecological specialisation and reproductive isolation.

### 1. Refining the history of divergence in the pea aphid complex

Our analyses reveal a reticulated evolutionary history in the pea aphid complex, reflecting high levels of shared polymorphism, incomplete lineage sorting, and ongoing or recent gene flow. The *Lathyrus pratensis* biotype consistently emerged as highly divergent, potentially representing a fully isolated lineage, consistent with the failure to find wild hybrids between this biotype and the other ones despite their occasional presence on *Lathyrus pratensis* plants [22]. Most other biotypes remain in a “grey zone” of speciation, with genome-wide *F*_ST_ values below 0.5, suggesting substantial gene flow persists [82].

Previous studies have reached contrasting conclusions regarding the timing of diversification among European pea aphid biotypes. Dating based on phylogenies of aphid bacterial endosymbionts suggested a recent origin (∼8,000 - 16,000 years ago) [83], although those estimates are likely strongly underestimated due to calibration using very recent introduction events in the USA. In contrast, dating based on mutation rates in asexual pea aphid generations and nuclear divergence between the *Lathyrus pratensis* and *Vicia cracca* biotypes suggested a much older origin (∼543,000 years [72]). Using an approach similar to [72] but with broader biotype sampling and accounting for sexual generations, we dated the split between *Lathyrus pratensis* and the rest of the complex back to a minimum of ∼545,000 years ago. Although this estimate may be biased by imperfect correction for ancestral polymorphism and by historical or ongoing gene flow, it suggests that the time required to complete reproductive isolation in pea aphids is comparable to what is reported in other insects [84,85]. By contrast, the most closely related biotypes (*Trifolium pratense* vs*. Onobrychis viciifolia*) appear to have diverged more recently, with a minimum estimate around 16,000 years ago, extending to ∼114,000 years ago for regions of the genome likely to act as old barriers (upper divergence time CI limit), although these values are probably underestimated due to gene flow. Together, these results suggest that divergence within the complex spans both ancient and more recent periods, with even the youngest splits predating Neolithic agriculture, challenging the hypothesis that reproductive isolation evolved rapidly in response to agricultural host-plant distributions [83]. Fazalova and Nevado (2020) [72] argued that the older age of the radiation leaves the possibility that pea aphids experienced numerous Pleistocene habitat fragmentations [86], during which they may have been confined to separate refugia [87] potentially inhabited by different host plants. This could imply phases of host plant specialisation and divergence in the absence of gene flow, such that the patterns of genetic differentiation and divergence – especially for distantly related biotypes – may reflect allopatric isolation followed by secondary contact, rather than divergence in the presence of gene flow.

Our analyses revealed strong, genome-wide correlations between *F*_ST_, *D*_XY_ and *π*, and their consistent evolution along the divergence continuum formed by the multiple biotype pairs, arguing against recurrent linked selection or ancient balancing selection as the primary drivers of differentiation. This contrasts with other well-studied systems in which recurrent (often linked) selection dominates genomic patterns (e.g., [30,35,68,88–91]). Instead, our results indicate heterogeneous gene flow among pea aphid biotypes shaping genomic landscapes of differentiation at all stages of divergence [35,67,68,92]. These results are in line with a smaller number of studies showing that divergent selection can be the primary determinant of genetic differentiation when ongoing gene flow constraints neutral divergence (e.g., in Darwin’s finches [92], monkeyflowers [34], *Castanopsis* trees [93]). By leveraging multiple biotype pairs spanning the divergence continuum, our study captures the gradual emergence of these genomic patterns in a way that has rarely been documented in natural systems. While our analyses support divergence with gene flow, we cannot fully exclude an initial phase of allopatric divergence followed by sufficiently ancient secondary contact for subsequent gene flow to dominate contemporary genomic landscapes of differentiation, and which may be detectable using demographic inference based on haplotype-resolved whole-genome data.

### 2. Genomic architecture of reproductive isolation along the continuum of divergence: pervasive role of reduced recombination and chromosomal rearrangements

Barrier loci were defined as 50-kb windows with significantly elevated *F*_ST_ and *D*_XY_, a combination expected for genomic regions resisting gene flow under a scenario of divergence with gene flow [67,68,73]. Demographic modelling that infer heterogeneous effective migration rates along the genome could provide complementary evidence for barrier loci when applied to individual-level genomic data [3,94]. However, several observations indicate that this operational definition provides a robust approximation of barrier loci in this system. First, the joint use of *F*_ST_ and *D*_XY_ is particularly appropriate in the pea aphid complex, where biotypes coexist in sympatry, since the correlation between these metrics across the divergence continuum is consistent with expectations under divergence with gene flow. Second, while elevated *D*_XY_ can also result from ancient polymorphism or mutation rate heterogeneity, the reduced *f*_dM_ observed in these regions indicates limited introgression, and their functional characteristics align with predictions derived from well-characterised barrier traits, providing strong arguments for their interpretation as barrier loci. Finally, although clusters of barrier loci may reflect linkage around a smaller number of causal loci particularly in rearranged or low-recombination regions, the persistence of numerous candidate regions even under stringent filtering suggests a highly polygenic architecture of reproductive isolation in pea aphids, even among closely related biotypes. This is consistent with previous phenotypic and QTL studies in the pea aphid, characterising several isolating traits (performance on host plants and host plant acceptance) whose polygenic bases are supported by available QTL studies [42,79].

Disentangling the causes of high differentiation remains particularly challenging in regions of low recombination rate, where linked selection can affect genetic differentiation and diversity [95–97]. Background selection typically elevates *F*_ST_ through reductions in within-population diversity but is not expected to systematically increase *D*_XY_ or reduce introgression. The combination of elevated *F*_ST_, elevated *D*_XY_, and reduced *f*_dM_ therefore argues against linked selection alone. Low recombination rates can also increase the variance of differentiation and diversity statistics, potentially inflating outlier detection in regions of low recombination [97]. Consistent with this, barrier loci were more often associated with low recombination rates, although recombination estimates based on Pool-Seq data may be biased by past selective or demographic events. Several lines of evidence nonetheless suggest this pattern does not solely result from linked selection (without gene flow): *F*_ST_ and *D*_XY_ did not exhibit higher variance in low-recombination regions compared to more recombining regions and inferred barrier loci were associated with reduced *f*_dM_. These observations are consistent with predictions that barrier loci are more likely to evolve and persist in regions of reduced recombination under gene flow [14,16,97–100], a scenario that pea aphid biotypes may have experienced for extended periods.

In agreement with theory, we also found that barrier loci tend to co-localise within chromosomal rearrangements. Although we did not resolve rearrangement breakpoints due to limitations of short-read Pool-Seq data, our results clearly show that a large rearranged region at the start of chromosome 1, comprising translocated and inverted segments, is strongly associated with reproductive isolation across all biotype pairs. The influence of this rearrangement is likely due to its large size and high density of barrier loci and outlier genes. Consistent with other empirical studies (reviewed in [102,103]), the pea aphid complex provides a clear example of chromosomal rearrangements acting as recombination modifiers, suppressing recombination in heterozygotes and increasing linkage disequilibrium among co-adapted alleles under gene flow (e.g., [18,102,103]).

We also observed tight physical linkage between outlier genes at smaller genomic scales, within and outside rearrangements. Although hitchhiking effects on nearby neutral loci could contribute to this pattern, we detected highly significant physical linkage among top outlier genes showing the strongest divergence and likely underlying reproductive isolation. Together, reduced recombination, chromosomal rearrangements, and physical linkage provide a favourable genetic architecture for the maintenance of linkage disequilibrium among regions resisting gene flow, thereby facilitating the coupling of barrier effects under ongoing gene flow [4,10–12,14,104–106].

Genome-wide *F*_ST_ and the number of barrier loci increase progressively with divergence from the *Trifolium* biotype, suggesting that reproductive isolation evolves through the gradual accumulation of small-effect mutations. However, the dynamics of genetic differentiation at barrier loci shows a different pattern that is consistent with threshold dynamics, characterised by alternating phases of rapid increase and stasis across the continuum. Although this interpretation of our results in terms of temporal dynamics should be taken with caution, the observed dynamics aligns with theoretical predictions of “tipping points”, i.e., sudden increase in RI due to genome hitchhiking and coupling processes [11,12,104,105,107–110]. In pea aphids, such dynamics may arise from reinforcement of host plant choice as a response to extrinsic selection against hybrids [12] and/or from changes in genetic architecture favouring clustering of barrier loci within chromosomal rearrangements [111]. Importantly, *F*_ST_ within barrier loci did not increase in a stepwise manner as net divergence increases when we excluded chromosomal rearrangements from the analysis (**S18D Fig**). The appearance of rearrangements and their segregation in the different biotypes may thus have had an important influence on the increase of divergence and/or the biotypes’ evolutionary histories.

Finally, we found no significant increase in the average length of barrier loci with divergence. The mechanism of “divergence hitchhiking” (in which strong divergent selection at a locus reduces effective gene flow over a large surrounding genomic region) would predict an increase in barrier loci length over time [112], although its overall importance remains debated [113–116]. While its signature has been reported in some systems (e.g., *Heliconius* butterflies [25]; *Populus* species [117]), its potential impact in pea aphid genomes seems limited or not fully measurable with our dataset. Instead, chromosomal rearrangements emerge as a stronger determinant of barrier accumulation. Based on a method applicable to Pool-Seq data, we found that biotypes in which the large rearrangement on chromosome 1 segregates at intermediate frequency show the highest proportion and largest barrier loci within this region. Although this contrasts with expectations that rearrangements should most strongly affect isolation when fixed [18], our observations may reflect the complex, nested nature of rearrangements, which likely vary in size, composition, and age among biotypes, as observed in other systems (*Timema cristinae* walking stick species [118]; *Mimulus* species [119]). Moreover, only subsets of these rearranged elements may be under divergent selection [14,18,108,111]. These results therefore call for a fine characterisation of the origin, structure, and evolutionary dynamics of these complex chromosomal rearrangements within and between pea aphid biotypes.

### 3. Nature of barriers to gene flow and their dynamics of recruitment along the continuum of divergence

We predicted that genes involved in host plant specialisation would be overrepresented among barrier loci, and focused on three functional categories: salivary effectors, chemosensory genes, and detoxification genes [60,61,78]. Salivary effector and detoxification genes are involved in nutritional pathways (phloem feeding), plant defense responses, and detoxification of secondary metabolites [120], potentially contributing to both pre-zygotic (selection against immigrants) and extrinsic post-zygotic isolation. In contrast, chemosensory genes, including chemoreceptors and odorant-binding proteins known for their roles in recognising compounds emitted by host plants and other sources [66,121,122], might be primarily involved in host plant acceptance/choice behaviour.

Candidate salivary effector genes were overrepresented mainly in “early” barrier loci (identified in closely-related biotype pairs, *F*_ST_ < 0.15), detoxification genes mainly in “late” barrier loci (identified only in the most divergent biotype pairs, *F*_ST_ > 0.25), and chemosensory genes were overrepresented in both. This suggests salivary effectors contribute early in divergence, chemosensory genes remain important throughout, and detoxification genes are more likely to act in later or intermediate stages. While different sets of chemosensory genes are identified as outlier genes in “early” versus “late” barrier loci, we did not find any evidence for the recruitment of different gene types (e.g., odorant versus gustatory receptors) at different stages of divergence. Overall, these results are consistent with phenotypic data showing that host plant adaptation traits have already evolved in the less divergent biotypes [22,39,41,43], and with theoretical expectations that host plant specialisation can evolve rapidly, partly via reinforcement, in a context of strong local adaptation in sympatry [123,124]. This classification as “early” or “late” barriers assumes that all biotype pairs follow a similar divergence trajectory. However, some barrier loci labeled as “late” could in fact have emerged early in the differentiation process but have evolved only in the most differentiated pairs. Additionally, our detection power might vary among biotype pairs, hence temporal interpretations should therefore be viewed cautiously. Nonetheless, these results provide insights into the relative contribution of these categories of genes at different levels of divergence, with host plant specialisation, hence ecological reproductive isolation, possibly intensifying as other genes from these functional categories, linked to existing or new specialisation phenotypes, come into play. Additionally, other categories of genes, directly involved in other forms of reproductive isolation or neutral with respect to reproductive isolation, tend to diverge later as a consequence of coupling/hitchhiking mechanisms, consistent with the observed “dilution” effect [11,12,105,106].

Focusing on the most differentiated outliers (“top” outlier genes), we identified 90 potential salivary effector genes and 16 chemosensory genes representing strong candidates underlying reproductive isolation between the *Trifolium* biotype and others. Fifteen salivary effectors and two chemosensory genes (Or3 and Or28) were top outliers in all four independent biotype pairs, suggesting shared functional determinants of host plant specialisation and recurrent targets of divergent selection, as seen in cichlid fish for opsin genes [29,125]. The substantial sharing of barrier loci across (independent) biotype pairs indicates parallel recruitment via introgression of alleles involved in ecological specialisation and reproductive isolation, or multiple adaptive mutations at the same loci.

Among top candidate salivary effector genes, we identified a sucrase gene (S1) previously linked to host plant specialisation in silkworms *Bombyx mori* [126]. Top chemosensory genes included several gustatory and odorant receptors, one ionotropic receptor, one OBP, and one CSP gene, often forming tightly linked clusters within (e.g., ORs) or across (e.g., ORs and OBP) multigene families. Close genomic proximity among top chemosensory genes may maintain linkage disequilibrium among loci of small effect on host acceptance, promoting coordinated evolution of host recognition traits. Detoxification genes were largely absent from top outliers (except for one GST detoxification gene acting also as a potential salivary effector), possibly due to limited detection power in biotype pairs with already high background differentiation. Most top candidate salivary effector and chemosensory genes have previously been reported as divergent among pea aphid biotypes (salivary effectors: [77]; chemosensory genes: [55–58]). The top chemosensory genes already identified include Or21, Or22 and Or3 within the large rearranged region on chromosome 1, Gr7-10 and Gr39 at the distal end of chromosome 1, and Or28 and CSP10 on chromosome 3. Strikingly, three QTL recently identified as controlling host plant acceptance in pea aphids [79] co-localise with two genomic regions highlighted here as prime candidates for reproductive isolation: the cluster of top chemosensory genes within the rearranged region on chromosome 1 and the location of Or28 on chromosome 3. These convergent results provide strong and independent support for the role of the regions we identified in host plant acceptance/choice, which could be the targets for future functional validation assays [127].

Top salivary effector and chemosensory genes were often physically clustered, particularly within the large rearranged region on chromosome 1 (30 salivary effectors and four chemosensory genes: OBP13, Or21, Or22, Or3) and at the distal end of chromosome 1 and around Or28 on chromosome 3. These patterns align with previous QTL mapping studies indicating co-localisation of host plant performance/survival and acceptance traits [42,50,51,79]. Specifically, our candidate region on chromosome 3 corresponds to a region of co-localised QTLs for host plant acceptance and survival previously identified [79]. Our results therefore provide evidence for concentrations of genes controlling host plant specialisation traits in at least three regions of the genome, consistent with the supergene model [128]. Similar patterns have been reported in other phytophagous insects, where host-use and preference traits map to the same genomic regions (*Timema cristinae* [129], *Rhagoletis pomonella* [130], *Spodoptera frugiperda* [132]). Close linkage between loci underlying traits under direct divergent selection and traits involved in assortative mating can be considered as “pseudo-magic” traits [6] or could mimic a single allele with pleiotropic effects [10], hence promoting divergence. In pea aphids, co-localisation of salivary effector and chemosensory genes within chromosomal rearrangements suggests that ecological specialisation is protected from gene flow by genetic coupling of host acceptance and survival traits, potentially facilitating their coordinated evolution under correlated environmental pressures [132]. Genetically linked outlier genes may act as stronger barriers to gene flow than isolated loci, and rearrangements may be positively selected to maintain advantageous allele combinations [115]. Comparative genomics across species within the *Acyrthosiphon* genus could clarify the evolutionary history, selection pressures, and role of these gene clusters in speciation [14,115].

In conclusion, our study provides new insights into the genomic landscape and functional nature of reproductive isolation at different levels of divergence in the pea aphid complex. Reproductive isolation emerges from a highly polygenic architecture shaped by heterogeneous recombination landscapes and chromosomal rearrangements that promote physical linkage among adaptive loci. The dynamic and functionally structured recruitment of genes involved in host plant recognition, interaction, and performance highlights how ecological pressures leave distinct genomic signatures at different stages of divergence. Together, these results illustrate how genomic architecture and ecological specialisation facilitate divergence with gene flow, providing a general framework for understanding the genetic basis of speciation in complex, polygenic systems.

## Material & methods

### 1. Resequencing data

In total, 13 European biotypes of pea aphids were chosen as representatives of the continuum of divergence within the pea aphid complex [22,40]. We used existing Pool-Seq data for 12 biotype populations (BioProject ID PRJNA385905), here named after their respective host plants, which were sampled in Eastern France from 2008 to 2012 [40,54,133]: *Trifolium pratense* (*n* = 60), *Onobrychis viciifolia* (*n* = 37)*, Medicago sativa* (*n* = 60), *Vicia cracca* (*n* = 29), *Medicago lupulina* (*n* = 27), *Lotus corniculatus* (*n* = 30), *Melilotus officinalis* (*n* = 34), *Pisum sativum* (*n* = 60), *Securigera varia* (*n* = 26), *Cytisus scoparius* (*n* = 29)*, Ononis spinosa* (*n* = 27) and *Genista tinctoria* (*n* = 30). All Pool-Seq sequencing libraries were prepared to ensure equal representation (equimolarity) of all individuals constituting each pool [54,133]. Guyomar et al. (2018) [133] also generated pool-Seq data for the biotype specialised on *Lathyrus pratensis*. However, this pool was found to contain migrants, i.e., individuals genetically assigned to other biotypes. As all pools were designed to avoid migrants, we built an artificial pool for this biotype by pooling reads of published *Lathyrus pratensis* individuals (10 individuals from United Kingdom [72] (PRJNA607096 and PRJNA607313) and three from France [134] (PRJNA255937). Before doing so we submitted all the individual data to a similar mapping and genotyping procedure (see next section) and performed a PCA to check that English and French samples grouped together (**S25 Fig**). We then down-sampled paired-end reads from all individuals to a read number corresponding to the lowest number of reads among the 13 individuals to create artificial equimolarity and we pooled all paired-end reads. This pool was then treated like the other ones in subsequent analyses.

As outgroup for *D*-statistics and *f*_dM_ estimation, we used individual resequencing data from two individuals of a Japanese population specialised on *Vicia cracca* (sample Sap05VC7, SRA: SRS2255006 and sample Iwamizawa, SRA: SRS2255007) from bioproject PRJNA255937). This population is known to have diverged before the European lineages [71], with the possible exception of the *Lathyrus pratensis* biotype (Jean Peccoud, personal communication), which we excluded from analyses requiring an outgroup. We pooled the two outgroup individuals following the same procedure as for the artificial *Lathyrus pratensis* pool.

### 2. Mapping and genotyping

We cleaned and trimmed all reads using *fastp* [135] with default parameters. We used *bwa-mem* [136] to map reads onto reference genomes combining the latest pea aphid chromosome level assembly from [63] called JIC1 (obtained from a clone of *Pisum sativum* biotype origin, available at https://bipaa.genouest.org/is/aphidbase/) and a set of genomes of endosymbionts frequently found in pea aphids (*Buchnera aphidicola* strain 5A (GCA_000021085.1), *Candidatus Fukatsuia symbiotica* strain5D (GCA_003122425.1), Candidatus *Hamiltonella defensa* strain A2C (GCA_003122445.1) and *Serratia symbiotica* (GCA_008370165.1)). We used *Picard* (https://broadinstitute.github.io/picard/) to filter out reads that mapped both onto the nuclear genome and onto the endosymbionts and mitochondrial genomes, and to mark duplicates. We called SNPs using the *bcftools* [137] mpileup function associated with the function bcftools call (parameters: -max-depth 250 -q 20 -Q 20). We then filtered the obtained vcf file using *vcftools* (parameters: --remove-indels --min-alleles 2 --max-alleles 2 --max-missing 0.9 --minQ 100 --minGQ 20 --minDP 10 --maxDP 200). We outputted both a VCF with only variable sites, as well as a VCF containing all called sites for the calculation of diversity-based statistics (*π* and *D_XY_*).

### 3. Continuum structure analysis

We used the *poolfstat* R package [138] to compute genome-wide pairwise *F*_ST_ averaged across all SNPs for all possible pairs of European biotypes and built a *F*_ST_-based dendrogram. We also used it to estimate the Patterson’s *D* statistics (a parsimony-like method to detect introgression) for all possible biotype quartet with the Japanese *Vicia cracca* biotype as outgroup, and *D*_min_, the minimum *D*-statistic for trios regardless of an assumed topology.

### 4. Estimation of summary statistics and recombination rate along the genome

We defined two sets of genomic windows for window-based analyses: non-overlapping 50-kb windows and overlapping 50-kb windows with a 10-kb overlap. Most estimations and analyses were done using non-overlapping windows, but we used overlapping windows for some analyses to increase the resolution where necessary.

#### Biotype-specific statistics

We estimated genetic diversity (*π*) along the genomes in non-overlapping 50-kb windows for all biotypes using the *npstat* software [139]. In pairwise analyses, we computed the genetic diversity of pairs as the average of the genetic diversity of the two biotypes of the pairs. We used *Relernn* [140] to estimate the recombination rate along the genome in the 13 pools. We set the mutation rate at 4.52×10^-10^ per generation per site, the generation time at 0.06667 years (i.e., a mutation rate of 6.78×10^-9^ per year per site), and the upper *rho* over theta ratio at 10. The mutation rate of 6.78×10^-9^ per year per site was computed by considering that pea aphids have on average 14 asexual generations per year with a mutation rate of 2.7×10^-10^ mutations per bp per generation (based on the asexual mutation accumulation experiment of Fazalova and Nevado 2020) and one sexual generation per year with a mutation rate of 3×10^-9^ mutations per site per generation (average of the estimates obtained in several insects [72]). *Relernn* outputs *rho* values for non-overlapping regions of an optimized (non-user chosen) window size along the genome. To obtain *rho* values in the non-overlapping 50k-b windows, we measured the weighted average of *rho* over overlaps between *Relernn* windows *versus* 50-kb windows. We also estimated gene density as the number of annotated genes overlapping the defined 50-kb windows.

#### Biotype pair-specific statistics

##### Biotype pair selection

In our “reference-based” approach, a biotype was used as the reference for comparison with the other 12 biotypes spanning a gradient of divergence. This design controls for genomic background and recombination landscape, provides a consistent reference for mapping divergence, and allows assessment of the dynamics of RI along the gradient since genetic distance from the reference can be quantified [30,31,37]. It therefore enabled a direct comparison of how genomic differentiation and divergence, introgression and barriers to gene flow vary across different levels of intraspecific divergence. We selected the *Trifolium pratense* biotype as the reference as it allowed us to build biotype pairs spanning the widest possible range of divergence (**Fig 1A**). We also defined a set of independent biotype pairs (i.e., with no shared biotype) for certain specific analyses. These pairs included one of the pairs from the previous set ( *Trifolium pratense vs Onobrychis viciifolia*) and three biotype pairs not including the *Trifolium pratense* biotype (*Medicago lupulina* vs*. Vicia cracca, Pisum sativum* vs*. Melilotus officinalis, Ononis spinosa* vs*. Genista tinctoria*). These four independent biotype pairs also span the continuum of divergence.

##### Divergence statistics

For all analyses using divergence statistics, we first filtered out windows containing fewer than 150 SNPs. Then, for each set of biotype pairs and for all retained genomic windows (both overlapping and non-overlapping windows), we estimated (1) relative divergence (i.e. genetic differentiation, *F*_ST_) among all pairs using *poolfstat* [138], (2) absolute divergence *D*_XY_ and (3) net divergence *D*_a_ statistics (only in non-overlapping windows) using a custom script (see jupyter notebooks provided as supplementary material) based on the VCF including non-variant sites. We converted *D*_a_ estimates into divergence time estimates using the computed mutation rate of 6.78×10 ^-9^ per year per site (see above).

##### Introgression statistics

We estimated the *f*_dM_ statistics [141] in 50-kb non-overlapping windows over the genome using the Pool-Seq estimator newly implemented in the *sliding.windows.fstat* function of the *poolfstat* package [138]. As detailed in Appendix A2 of the package vignette v3.1 [142], the estimator leverages on an equivalent (re)formulation of the numerator and denominator of *f*_dM_ in terms of F4 and F3 statistics, which enables straightforward derivation of unbiased estimators for Pool-Seq data (especially for the denominator of the statistics). The *f*_dM_ was computed for eight out of 12 biotype pairs, with the aim of quantifying the level of introgression between the *Trifolium* biotype and the second biotype in the pair (representing P2 and P3). This statistic quantifies the shared variation between P3 and P2 or between P3 and P1 of the example topology presented in **S26 Fig** [70]. *f*_dM_ gives positive values for introgression between P3 and P2 and negative values for introgression between P3 and P1. We thus composed quartets of biotypes with *Trifolium* and the tested biotype as P2 or P3, the Japanese *Vicia cracca* outgroup as P4, and another biotype as P1. To identify the most suitable P1 biotype to use, we computed the *D*-statistic using different biotypes as P1 and chose those yielding positive values. For the biotype pairs *Trifolium pratense - Medicago sativa* and *Trifolium pratense - Medicago lupulina*, no such biotype could be found, so we did not compute *f*_dM_ for those pairs. We could not compute *f*_dM_ for the closest biotype pair *Trifolium pratense - Onobrychis viciifolia*.

Finally, we did not compute *f*_dM_ for the pair *Trifolium pratense - Lathyrus pratensis* as the latter may have diverged before the Japanese *Vicia cracca* biotype. All analyses including *f*_dM_ were restricted to windows with positive values, where the *f*_dM_ statistic indicated introgression between P3 and P2, i.e., between the tested biotype and *Trifolium pratense* (**S26 Fig**).

#### Correlation among genome-wide statistics

We estimated genome-wide correlations between the statistics mentioned above using non-overlapping 50-kb windows via Spearman correlation tests in R version 3.6.1. We fitted linear, asymptotic and hyperbolic (Michaelis–Menten) models using the nls() and lm() functions in R (v 4.5.1, stats package). The hyperbolic model was tested for the relationship between *f_dM_*and *D_a_*, as it allows a fitted maximum (observed *f_dM_*) and asymptotes to zero at high divergence.

### 5. Barrier loci identification and analysis

We defined barrier loci as 50-kb windows with both significantly elevated *F*_ST_ and significantly elevated *D*_XY_ [67,68]. To assess significance levels, we followed the permutation test procedure and scripts used in [69]. Briefly, we Z-transformed the *F*_ST_ and *D*_XY_ values of all genomic windows based on their mean and standard deviations in the three autosomes and the X chromosome separately, and each biotype pair independently. We then performed permutations of the genomic windows and applied a Savitzky-Golay smoothing filter to the shuffled Z-transformed values, using a polynomial order of 3 fitted to seven windows [143] following [69] scripts. For each window, we compared the original Z-transformed values to the distribution of smoothed values across 10,000 permutations, and calculated the *p*-values based on the number of permutations with Z-transformed values as large or larger than the original value. We then selected windows as barrier loci when both *F*_ST_ and *D*_XY_ were associated with *p*-values < 0.05. All other windows with non-significant *F*_ST_ or *D*_XY_ estimates were considered as non-barrier loci. We did not apply false discovery rate (FDR) correction in this procedure because barrier loci are expected to involve many loci of small effect, such that strict multiple-testing correction would bias detection towards only the strongest peaks [144–146]. Moreover, significance was assessed using permutation-based genome-wide null distributions for each biotype pair, which already limit false positives, and barrier loci were defined based on the joint significance of *F*_ST_ and *D*_XY_, consistent with recommendations to combine multiple statistics to reduce false discoveries in genome scans [147,148]. Finally, we eventually combined several lines of evidence, by testing whether identified barrier loci also showed reduced introgression rate (*f*_dM_). Nevertheless, to assess the robustness of our results, we repeated the main analyses using Benjamini–Hochberg FDR–corrected *p*-values; although this reduced the number of detected barrier loci, the overall patterns and conclusions remained unchanged ( **S27 Fig**). We applied this procedure separately to the non-overlapping and overlapping genomic window sets. Most barrier loci analyses were done using non-overlapping windows, with overlapping windows being used to increase the resolution when merging adjacent barrier loci. This merging allowed us to estimate the total number of barrier regions,the average length of barrier regions and mapping annotated genes onto barrier loci.

To test whether barrier loci were non-randomly concentrated in specific genomic regions in each biotype pair, we performed a permutation-based enrichment analysis. The genome was partitioned into consecutive 15-Mb bins within each chromosome, and we counted the observed number of barrier loci falling within each 15-Mb bin. To assess statistical significance, we generated a null distribution by randomly reallocating the same number of barrier loci across all available genomic windows for that biotype pair, while preserving chromosome structure (*n* = 10,000 permutations). For each bin, an empirical *p*-value was computed as the proportion of permutations in which the number of barrier loci was greater than or equal to the observed count. To account for multiple testing across genomic bins, empirical *p*-values were adjusted using the Benjamini–Hochberg false discovery rate (FDR) correction. Bins with an FDR-adjusted *p*-value below 0.05 were considered significantly enriched in barrier loci. We further quantified overlap among biotype pairs by counting, for each genomic bin, the number of biotype pairs showing significant enrichment.

We assessed the dynamics of barrier loci as net divergence (*D*_a_) increases across the continuum of divergence by assessing how (i) the total base pairs within barrier loci, (ii) the total number of barrier regions (defined as merged adjacent or overlapping barrier loci), (iii) the average length of barrier regions and (iv) the average *F*_ST_ within barrier loci varied as a function of divergence with the *Trifolium* biotype (measured by *D*_a_). For the analysis of *F*_ST_, we used cubic regression splines (CRS, *crs* package in R). Models were fit with *D*_a_ as the response and *F*_ST_ as the predictor, using exhaustive cross-validation to determine optimal number of segments and knots. Spline degree and number of segments were recorded, and model fit was assessed via residual sum of squares (RSS) and cross-validation (CV) score. Barrier and non-barrier loci were analysed separately to compare the abruptness of changes in *F*_ST_ between the two datasets. We also performed Wilcoxon rank sum tests to compare the recombination rate and *f*_dM_ within barrier loci versus within non-barrier loci.

### 6. Analysis of chromosomal rearrangements

To establish chromosomal rearrangements, we aligned the four chromosomes of the *A. pisum* assembly generated from an individual specialised on *Medicago sativa* (AL4f) (GCA_005508785.2, [74]) and the four scaffolds of the *A. pisum* assembly generated from an individual putatively specialised on *Pisum sativum* (JIC1) [63]. Dot plots allowed us to identify approximate boundaries of several candidate chromosomal rearrangements between those two genomes. We then performed Fisher’s exact tests to investigate the overrepresentation of barrier loci within those chromosomal rearrangements.

We evaluated the segregating frequency of a large rearrangement located in the first ∼39Mb of chromosome 1 (scaffold 2 in the JIC1 version of the genome) in each biotype, following a method recently proposed for Pool-Seq data [75]. Briefly, we considered that two given biotypes that have been exchanging genes should present a good correlation of allele frequencies, except within a chromosomal rearrangement (potentially an inversion) segregating at different frequencies between biotypes. Within the rearrangement (which contains outlier loci), some alleles could be (nearly) differentially fixed. This situation allows inferring the allele linked to either variant of the rearrangement, hence reconstructing haplotypes. Allele frequencies at these SNPs thus provide a reliable estimate of the rearrangement frequency in each population [75]. We identified such “diagnostic” SNPs as those with the highest *F*_ST_ (>80% quantile of the *F*_ST_ distribution) within the first 39 Mb of chromosome 1 in at least 8 out of the 12 biotype pairs (each including the biotype on *Trifolium pratense*).

Finally, we repeated the analyses assessing the dynamics of barrier loci as net divergence increases along the continuum of divergence when excluding potential rearranged regions to test the effect of chromosomal rearrangements on these patterns.

### 7. Functional gene content analysis

We defined outlier genes as those overlapping with merged overlapping barrier windows in any biotype pairs. The pea aphid genome has benefited from high-quality annotation including manual annotation of different categories of genes on the first assembly version [64,65,80] but we realised during this study that some annotation information has not been carried through the later assembly versions and newly produced assemblies, including the reference genome we used here, JIC1. As a consequence, we could not rely on a good-quality functional annotation at the genome scale, preventing us from performing Gene Ontology enrichment searches without any prior hypothesis. Instead, we first generated our own annotation file by transposing the annotation of the first pea aphid genome assembly [80] onto the JIC1 genome (**S5 Table**). Then, we focused on gene categories (chemosensory genes, potential salivary effector genes and detoxification genes) for which we could retrieve high-quality annotations and which were expected to be involved in host plant specialisation and reproductive isolation [60,78].

For chemosensory genes, we combined previous annotations [55,58,64,66] to obtain a list of 199 genes classified as “OR” (olfactory receptors), “GR” (gustatory receptors), “IR” (Ionotropic Receptors), “SNMP” (sensory neuron membrane proteins which have been shown to be a signalling component required for pheromone sensitivity in *Drosophila* [149]), “CSP” (chemosensory proteins) or “OBP” (odorant binding protein involved in the transport of odorants) (**S5 Table**). These genes have been annotated on two previous versions of the pea aphid genome (Acyr_2.0 – GCF_000142985.2 (The International Aphid Genome Consortium 2010) and GCA_005508785.2 [74]), so we converted their coordinates into those of the JIC1 assembly. For candidate salivary effectors, we first recovered the gene IDs of 3546 transcripts expressed in pea aphid salivary glands and with a secretion signal according to [59] as well as 1,485 transcripts whose expression is up regulated (compared to the alimentary tract) in either or both the *Pisum sativum* biotype and the *Medicago sativa* biotype. Most of those transcripts were annotated on previous versions of the pea aphid genome, so we converted their coordinates into those of the JIC1 genome. We eventually recovered 2,957 potential salivary effectors (**S5 Table**). For detoxification genes, we recovered 94 genes annotated as being similar to cytochrome P450 proteins (CYP450 enzymes known to play key roles in the detoxification of plants secondary metabolites [61]), as well as glutathione S-transferases (GSTs) and Uridine diphosphate (UDP)-glycosyltransferases, which detoxify metabolites by conjugating them with polar molecules [61] (**S5 Table**). To evaluate the overrepresentation of each gene category among outliers, we performed Fisher’s exact tests across all biotype pairs combined (i.e., all outlier genes) and for each biotype pair separately.

To analyse the nature of outlier genes (those within barrier loci) along the continuum of divergence, we first classified barrier loci as “early” *versus* “late”. “Early” barrier loci were those considered as barrier loci in at least two biotype pairs among the four pairs with lowest global *F*_ST_ (i.e., *Trifolium pratense* vs. {*Onobrychis viciifolia*; *Medicago sativa*; *Medicago lupulina*; *Vicia cracca*}); “late” barrier loci were those considered as barrier loci in at least two biotype pairs among the four pairs with highest global *F*_ST_ (i.e., *Trifolium pratense* vs. {*Cytisus scoparius*; *Ononis spinosa*; *Genista tinctoria*; *Lathyrus pratensis*) but not identified as barrier loci in any of the four biotype pairs with lowest *F*_ST_. Note that this classification implies that some barrier loci labelled as “late” could in fact have emerged early in the divergence process but have evolved uniquely in biotypes that diverged first. We then tested the overrepresentation of genes from each functional category within “early” or “late” barrier loci (separately), using Fisher’s exact tests.

We then narrowed down the list of outlier genes to the most promising ones for further analyses. To this aim, we first classified, for each biotype pair, outlier genes into three categories according to the significance of the barrier loci in which they occurred, requiring concordant statistical support from both *F*_ST_ and *D*_XY._ Outlier genes were considered as “significant” when both statistics were associated with *p*-values < 0.05, “highly significant” when both had *p*-values < 0.01, and “very highly significant” when both had *p*-values < 0.001. We then defined “top” outlier genes as the ones most significantly and repeatedly identified as outlier across all biotype pairs involving the *Trifolium* biotype. Our criteria required that the gene be outlier in at least seven biotype pairs and detected as “highly” or “very highly” significant outlier gene in at least five biotype pairs. We then analysed this list of “top” outlier genes by assessing their nature and distribution along the genome ( **Fig 6**). Physical clustering of top outlier genes was quantified by calculating nearest-neighbour (NN) distances between consecutive top outlier genes along each chromosome, accounting for annotated gene positions and gene density. For each chromosome, the percentage of top outlier genes whose nearest neighbour was located within 10 kb (parameter X_kb = 10) was computed as a statistic for physical clustering. To evaluate the significance of clustering, we performed a permutation test (*n* = 10,000). In each iteration, the same number of genes as observed outliers was randomly sampled from the pool of all annotated genes on the corresponding chromosome (either all genes or only genes outside candidate chromosomal rearrangements, depending on the analysis). The same NN distance metric was calculated for these randomly sampled genes, and the mean percentage within 10 kb across chromosomes was recorded. Empirical *p*-values were calculated as the proportion of permutations in which the simulated statistic was equal to or higher than the observed percentage. Distributions of observed NN distances and permutation test results were visualised in histograms. Analyses were performed separately for (i) all top outlier genes and (ii) top outlier genes located outside candidate chromosomal rearrangements.

We finally identified outlier genes potentially involved in the divergence of several independent biotype pairs and representing shared reproductive isolation genes and/or generic functional determinants of host plant specialisation. We did so by identifying “top” outlier genes in the four independent biotype pairs, i.e., genes consistently detected as outliers in all four pairs and showing the strongest signals of divergence in these pairs (a gene must be “highly” or “very highly*”* significant in at least three out of four of these independent biotype pairs).

Code developed to perform the analyses is available as supporting information (notebook_all_analyses.html).

## Acknowledgments

This study was supported by the funded projects ANR SPECIAPHID N° ANR-11-BSV7-0005 and ANR MECADAPT N° ANR-18-CE02-0012-03. We thank Thibault Leroy, Alan Le Mohan, Anja Westram, Charles Perrier, Simon Boitard and Jules Romieu for methodological reflections and suggestions on how to analyse this complex dataset. We thank Roger Butlin for comments and suggestions on an earlier version of the manuscript.

## Authors contributions

MR: conceptualisation, methodology, formal analysis, writing: original draft; JJ: contribution to data acquisition and preliminary data analysis steps, writing: review and editing; JP: contribution to data acquisition, preliminary data analysis steps, writing: review and editing; LM: contribution to data acquisition and preliminary data analysis steps; FM: resources: preparation of the Pool-Seq sample of the *Cytisus* biotype; FL: resources: support for access to HPC Genouest server, lifting of gene annotations onto JIC1 reference genome; RV: funding acquisition, methodology, writing: review and editing; EJ: funding acquisition, methodology, writing: review and editing; JCS: funding acquisition, resources: coordination of sample and data acquisition; writing: review and editing; MG: funding acquisition, methodology, method development, writing: review and editing; CMS: funding acquisition, study coordination, conceptualisation, supervision, methodology, formal analysis, writing: original draft. All authors proofread and approved the final version of the manuscript.

## Supporting Information

### Supplementary tables

S1 Table. Correlations with and without *Lathyrus* biotype pair.

S2 Table. Statistics associated with all 50kb genomic windows.

Tab1: non-overlapping windows; tab2: overlapping windows.

S3 Table. Barrier enrichment results. S4 Table. Outlier genes.

S5 Table. Curated gene annotation on JIC1 reference genome.

### Supplementary figures

supplementary_figures.pdf

S1 Fig. Genome-wide relationships between *F*_ST_ and *D*_XY_ across the continuum.

S2 Fig. Genome-wide relationships between genetic diversity (*π*) and *D*_XY_ across the continuum. Red dotted lines represent identity lines.

S3 Fig. Genome-wide correlations between divergence and diversity statistics in reference-free independent pairs. Correlations between *F*_ST_, *D*_XY_ and *π* in four independent biotype pairs of increasing divergence. In each plot, dots represent the different genomic windows for which the different statistics could be computed, with two focal statistics plotted on the x- and y-axes and the third statistic represented by the dot colour code. Grey dotted lines represent linear regression fits, and red dotted lines represent identity lines.

S4 Fig. Dynamics of the strengths of correlations between diversity and divergence statistics as net divergence increases in reference-free independent pairs

S5 Fig. Genome-wide relationships between genetic diversity (*π*) and *F*_ST_ across the continuum.

S6 Fig. Genome-wide relationships between gene density and genetic diversity (*π*) across the continuum.

S7 Fig. Genome-wide relationships between gene density and *D*_XY_ across the continuum.

S8 Fig. Genome-wide relationships between gene density and *F*_ST_ across the continuum.

S9 Fig. Genome-wide relationships between recombination rate and genetic diversity (*π*) across the continuum.

S10 Fig. Genome-wide relationships between recombination rate and *D*_XY_ across the continuum.

S11 Fig. Genome-wide relationships between recombination rate and *F*_ST_ across the continuum.

S12 Fig. Genome-wide relationships between *F*_ST_ and *f*_dM_ across the continuum. This shows only positive *f*_dM_ values.

S13 Fig. Genome-wide relationships between *D*_XY_ and *f*_dM_ across the continuum. This shows only positive *f*_dM_ values.

S14 Fig. Evolution of the strength of Spearman correlations between *f*_dM_ and both *F*_ST_ (left) and *D*_XY_ (right) as divergence increases along the continuum. Colours correspond to biotypes in Figure 1A.

S15 Fig. Distribution of *f*_dM_ within and outside of inferred barrier loci. This shows only positive *f*_dM_ values.

S16 Fig. Manhattan plots of the landscapes of *F*_ST_ (top), *D*_XY_ (bottom) and the localisation of inferred barrier loci (green dots) and shared barrier across all pairs (red dots) in the four independent biotype pairs of increasing divergence (top to bottom).

S17 Fig. Number of unique versus shared barrier loci among the four independent pairs of biotypes. All categories are mutually exclusive.

S18 Fig. Dynamics of genomic regions acting as barriers to gene flow as divergence increases across the continuum, when candidate chromosomal rearrangements are excluded from the analyses. Coloured dots represent biotype pairs as in Figure 1. **A**: Total base pairs in barrier regions (i.e., merged adjacent barrier loci) loci as net divergence (*D*_a_) increases (in Mb), **B**: Number of (merged) barrier regions as net divergence increases, **C**: Average length of (merged) barrier regions as net divergence increases, **D**: Mean *F*_ST_ within barrier windows as net divergence increase (opaque points) and mean *F*_ST_ in non-barrier windows (randomly chosen among all non-barrier windows to obtain the same number of non-barrier windows as the number of identified barrier windows, for each biotype pair) as divergence increases (transparent points). Red (for barrier loci) and black (for non-barrier loci) dashed lines show regression spline (CRS) fits; orange (for barrier loci) and grey (for non-barrier loci) dashed lines show asymptotic fits.

S19 Fig. Relation between chromosome 1 structural rearrangement segregating frequency and the average size of barrier loci within it.

S20 Fig. Dot plots of chromosomes A2, A3 and X from the *A. pisum* assembly based on a *Medicago sativa* individual AL4f (GCA_005508785.2) (Li et al. 2019) *versus* scaffolds 3, 4 and 1, respectively, from the *A. pisum* assembly based on a putative *Pisum sativum* individual JIC1 (Mathers et al. 2021).

S21 Fig. Distribution of recombination rate within and outside of inferred barrier loci in windows of low (left) *versus* high gene density (right). Significant Wilcoxon rank sum tests comparing the recombination rate within barrier loci *versus* within non-barrier loci are indicated with a star.

S22 Fig. Dispersion of *F*_ST_ and *D*_XY_ values in bins of low, middle and high recombination rate in three biotype pairs: A: *Medicago sativa - Trifolium*, B: *Pisum sativum - Trifolium*, and C: *Lathyrus pratensis - Trifolium*.

S23 Fig. Enrichment tests of chemosensory (A), salivary effectors (B) and detoxification (C) genes within barrier loci across the continuum. Biotypes are sorted by increasing divergence with the *Trifolium* biotype.

S24 Fig. Physical clustering of top outlier genes. Panels A and C show the distribution of nearest-neighbour (NN) distances among top outlier genes, with grey bars representing counts of genes in each 5 kb bin, and red vertical lines indicating the observed NN statistic. Panels B and D show the distribution of permutation statistics across 10,000 randomisations, with grey bars representing the random expectation and the red vertical line showing the observed statistic. Panels A–B include all top outliers, while C–D include only top outliers outside candidate chromosomal rearrangements. The 10 kb threshold indicates the a priori definition of “close” genes.

S25 Fig. PCA of individual *Lathyrus pratensis* samples subsequently pooled *versus* pooled samples of the rest of the biotypes.

S26 Fig. Schematic of the reasoning of the computation of the *f*_dM_ statistic in the context of the continuum.

S27 Fig. Reproduction of the main results when applying FDR correction on *p*-values for *F*_ST_ and *D*_XY_ before identifying barrier loci. (1) Equivalent to Figure 3; (2) Equivalent to Figure 4; (3) Functional enrichment results for the three candidate gene categories; (4) Equivalent to Figure 6.

## Supplementary figures

**Figure S1:**
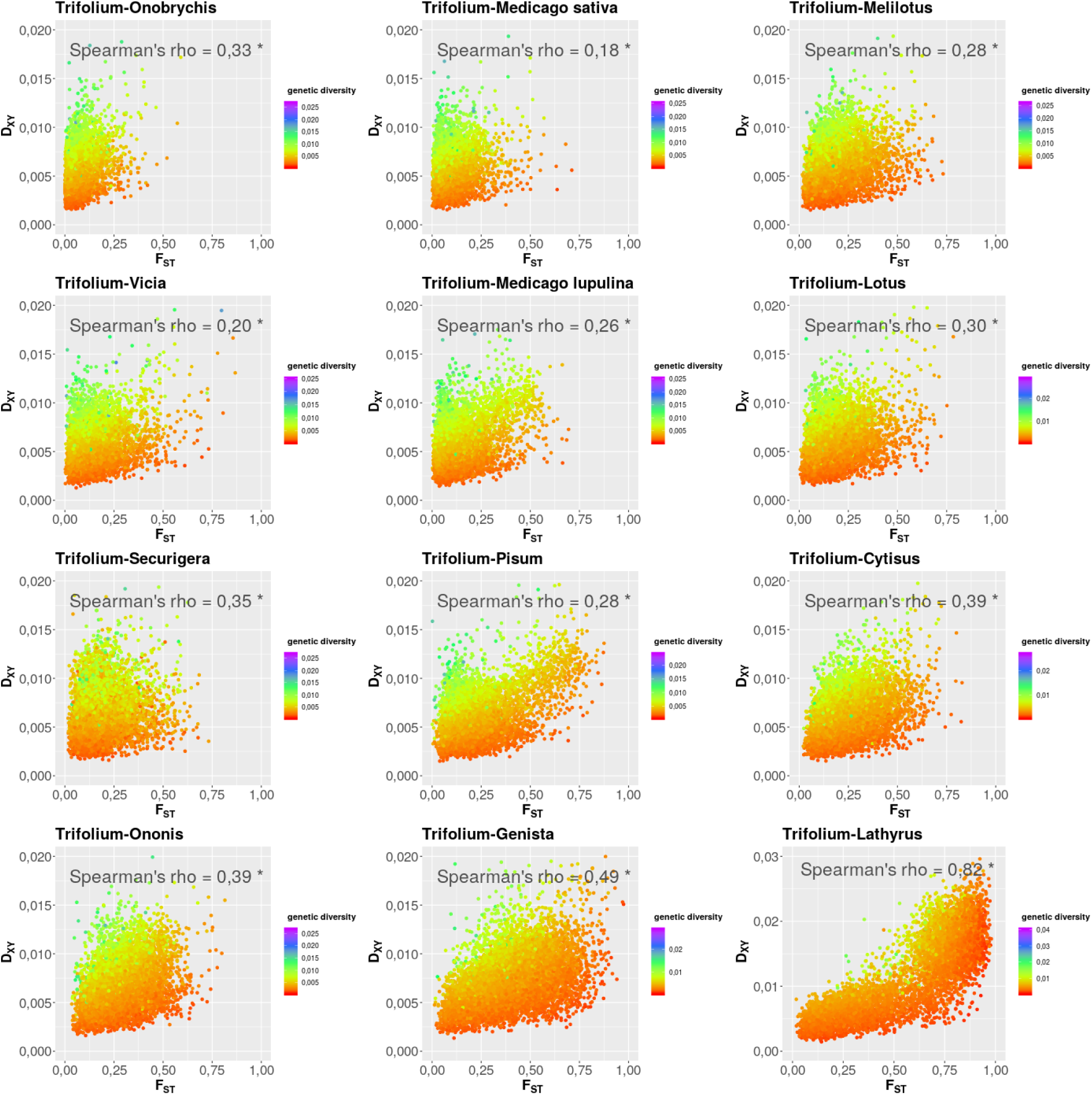
Genome-wide relationships between *F_ST_* and *D_XY_* across the continuum.

**Figure S2:**
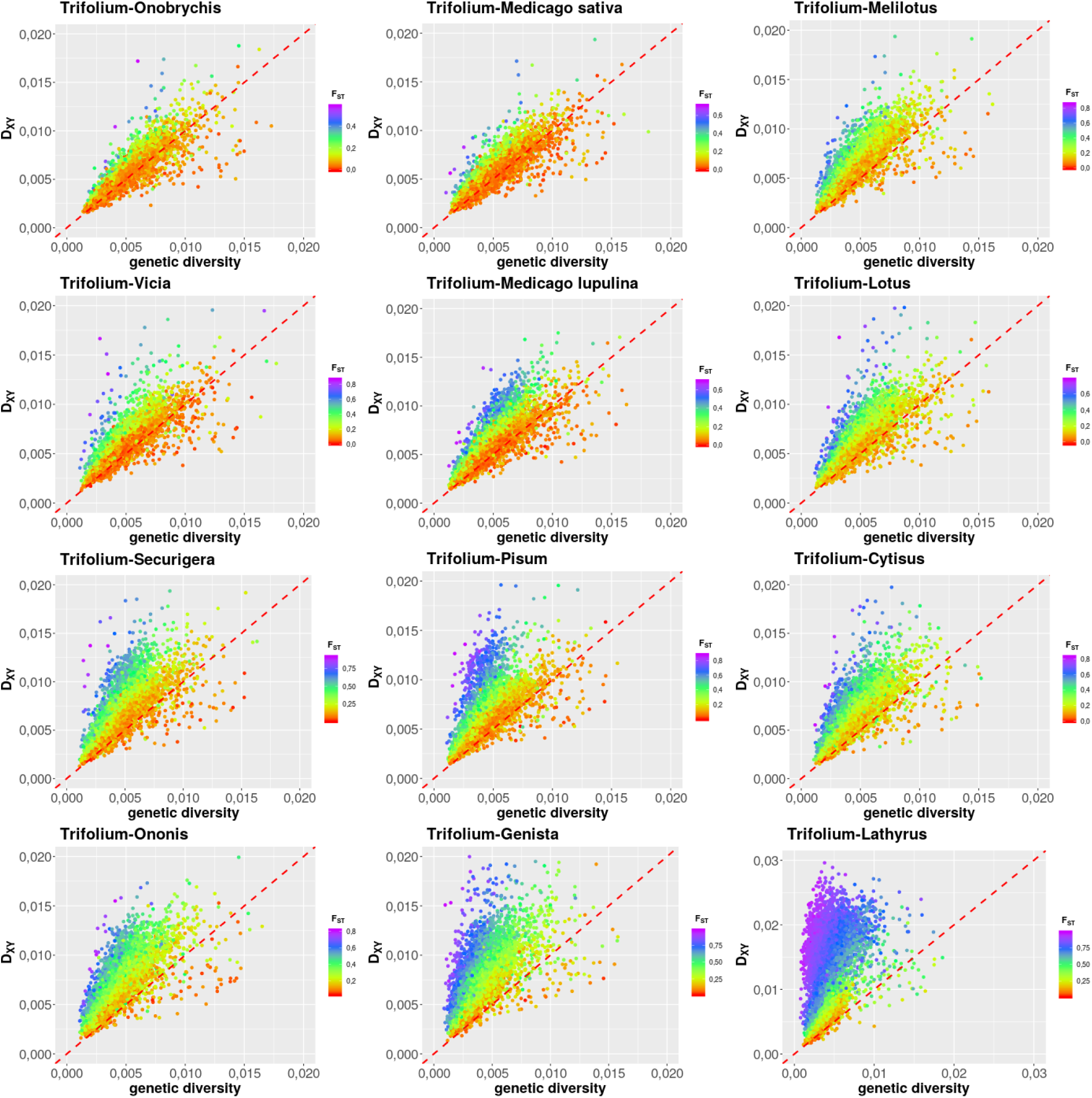
Genome-wide relationships between genetic diversity (*π*) and *D_XY_* across the continuum. Red dotted lines represent identity lines.

**Figure S3:**
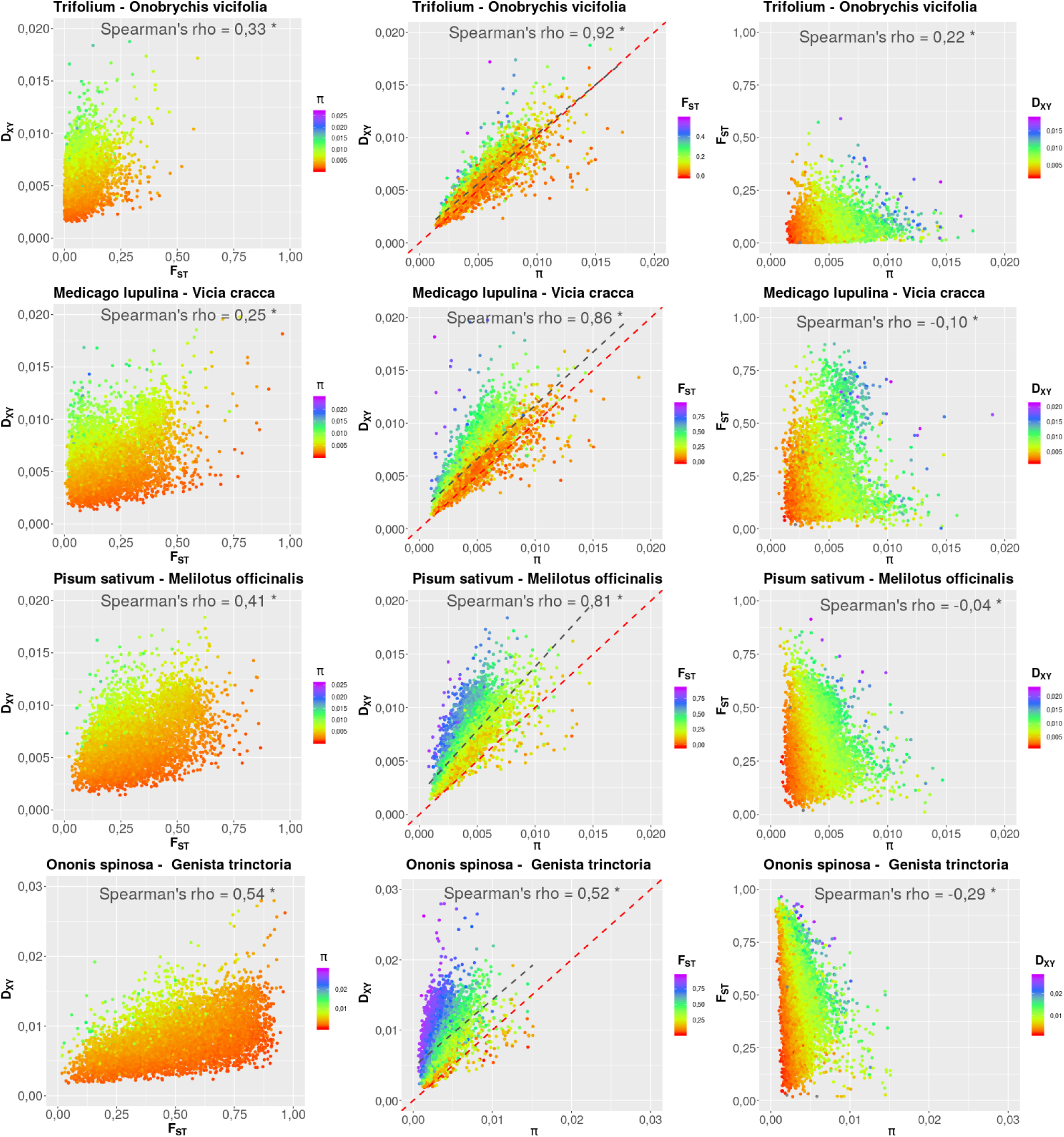
Genome-wide correlations between divergence and diversity statistics in reference-free independent pairs. Correlations between *F_ST_*, *D_XY_* and *π* in four independent biotype pairs of increasing divergence. In each plot, dots represent the different genomic windows for which the different statistics could be computed, with two focal statistics plotted on the x- and y-axes and the third statistic represented by the dot colour code. Grey dotted lines represent linear regression fits, and red dotted lines represent identity lines.

**Figure S4:**
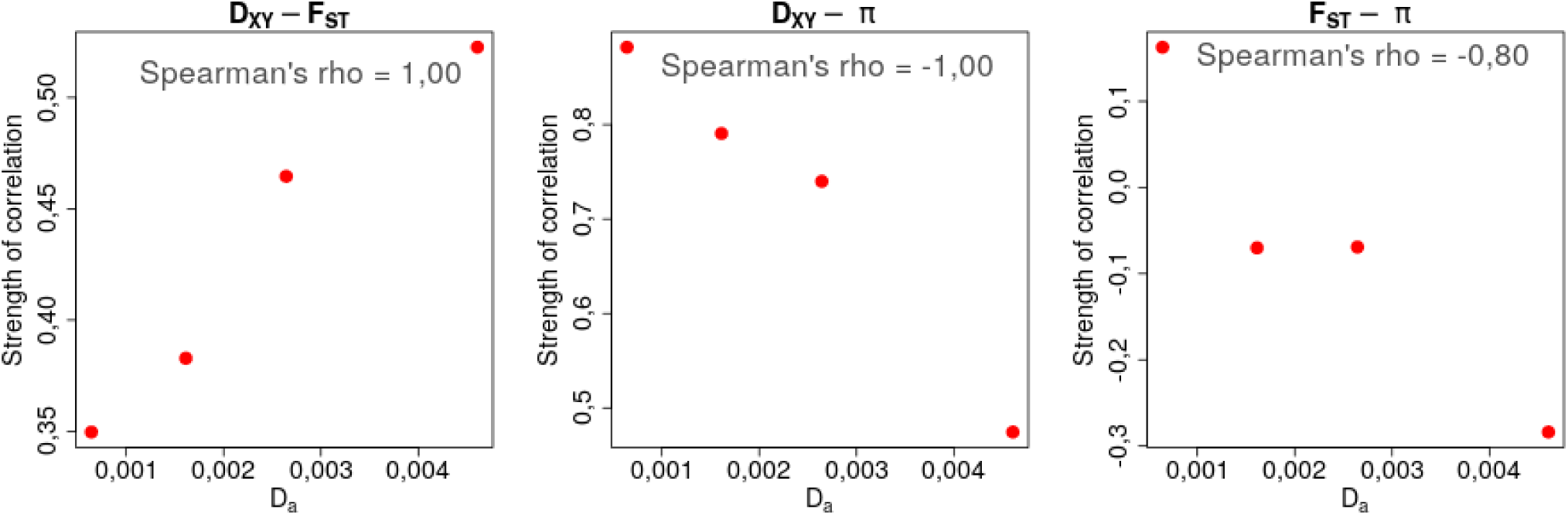
Dynamics of the strengths of correlations between diversity and divergence statistics as net divergence increases in reference-free independent pairs.

**Figure S5:**
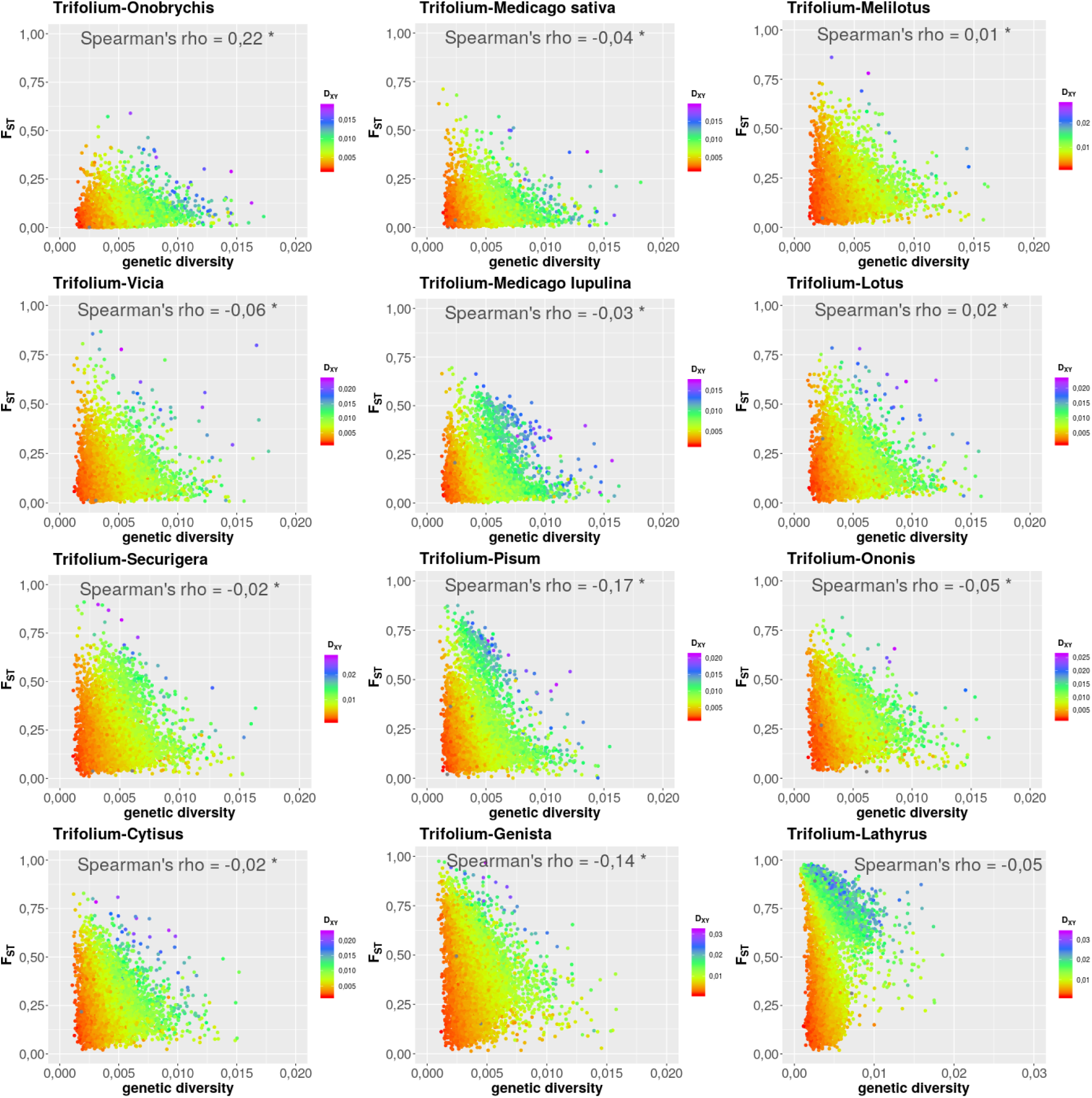
Genome-wide relationships between genetic diversity (*π*) and *F_ST_* across the continuum.

**Figure S6:**
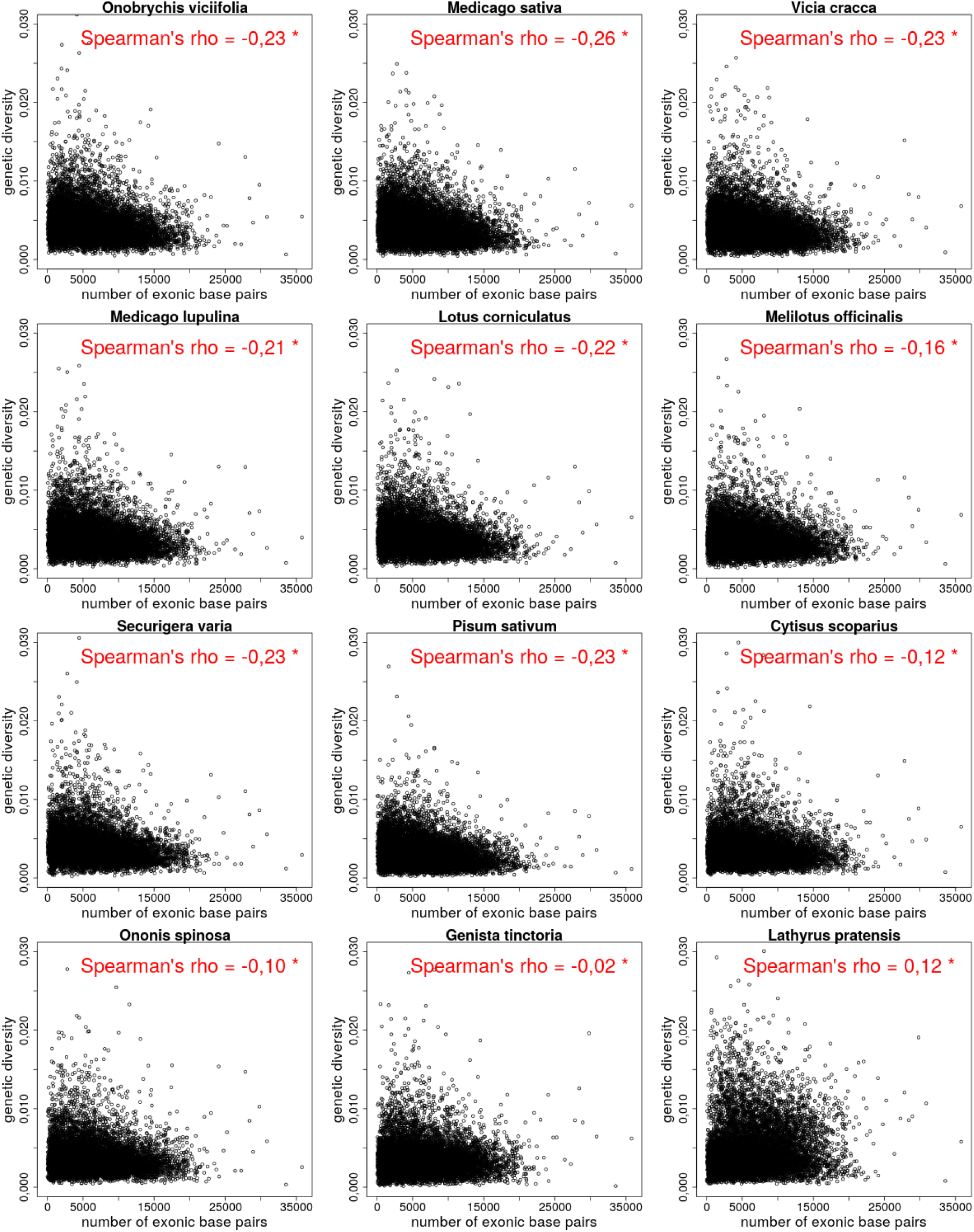
Genome-wide relationships between gene density and genetic diversity (*π*) across the continuum.

**Figure S7:**
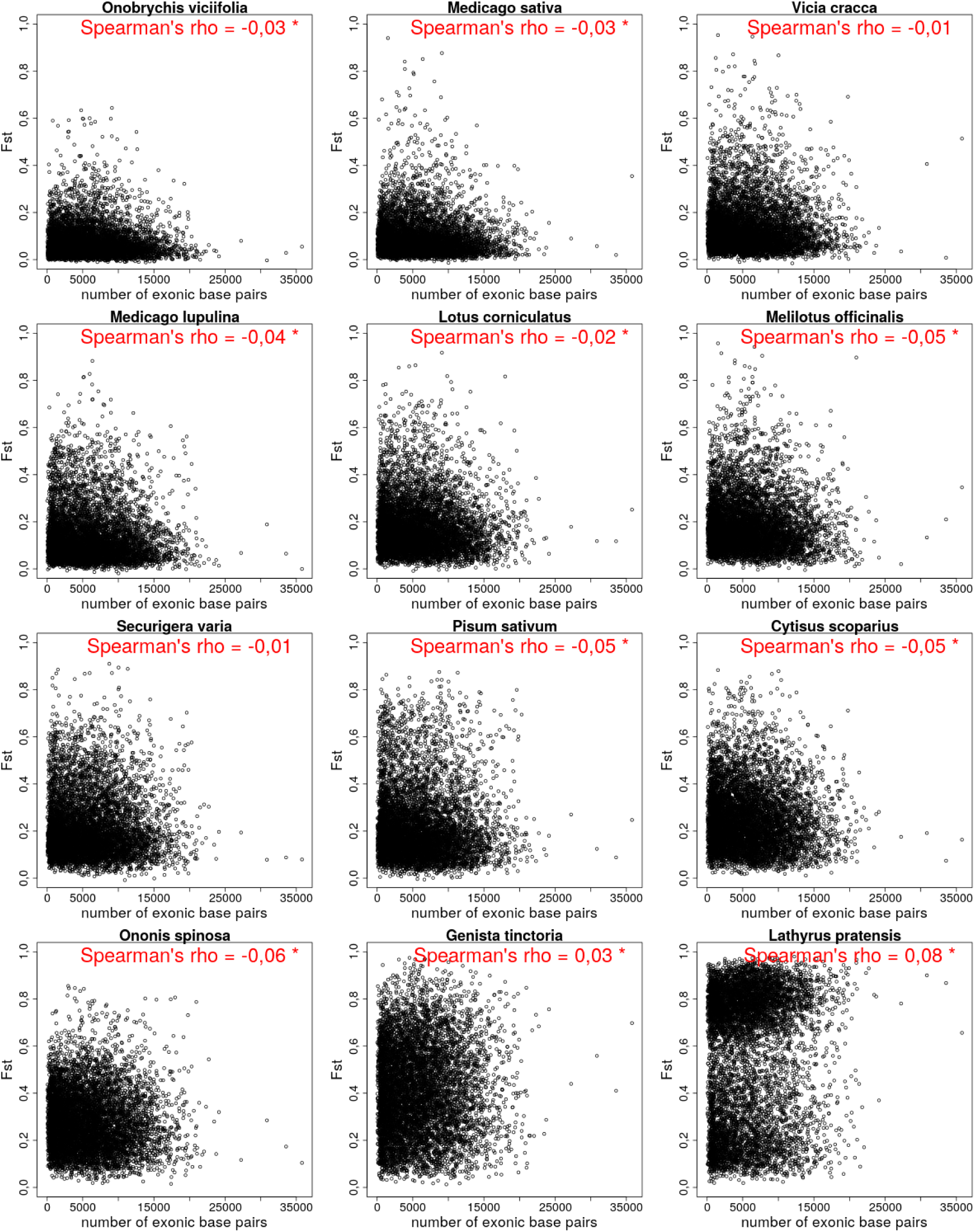
Genome-wide relationships between gene density and *D_XY_* across the continuum.

**Figure S8:**
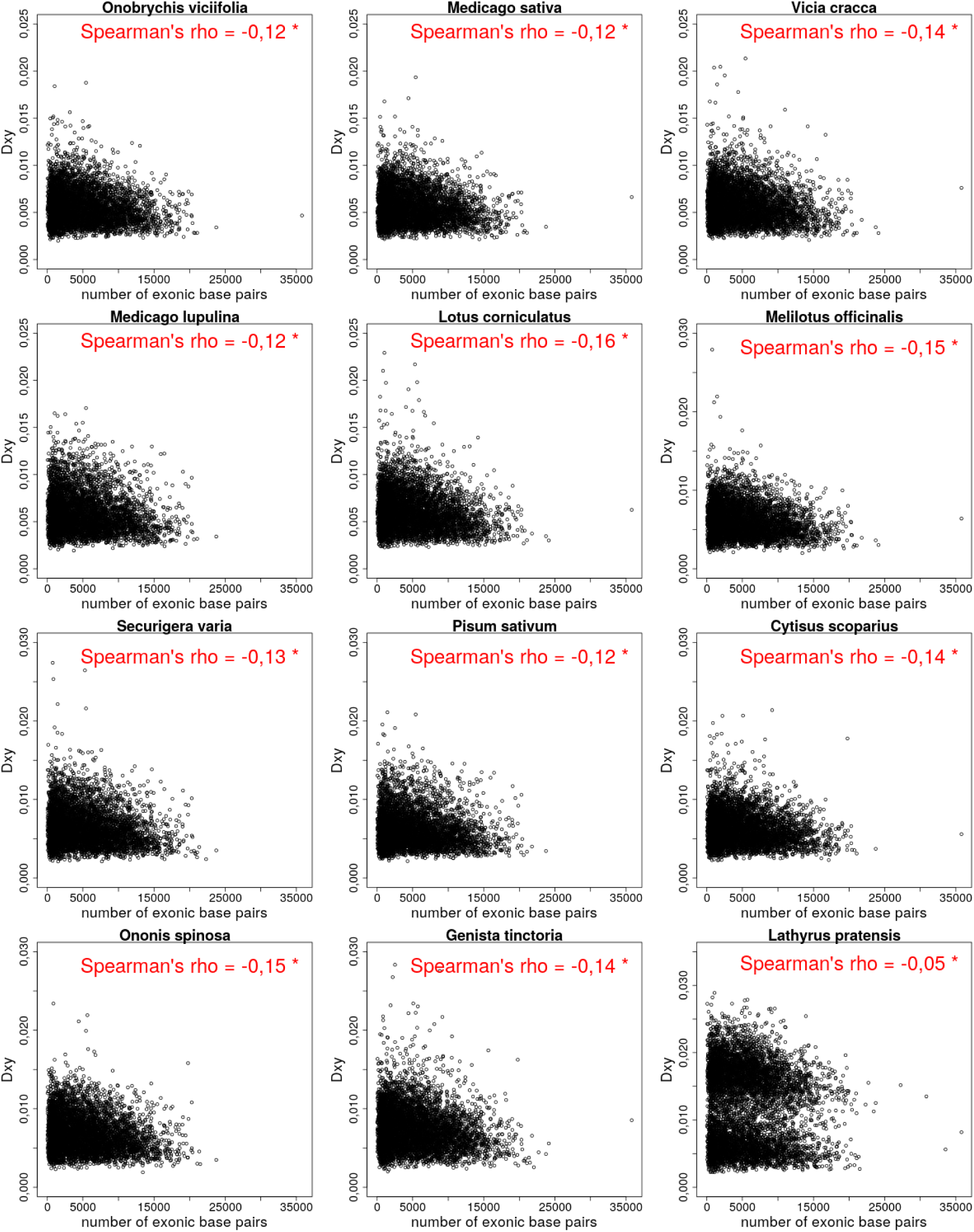
Genome-wide relationships between gene density and *F_ST_* across the continuum.

**Figure S9:**
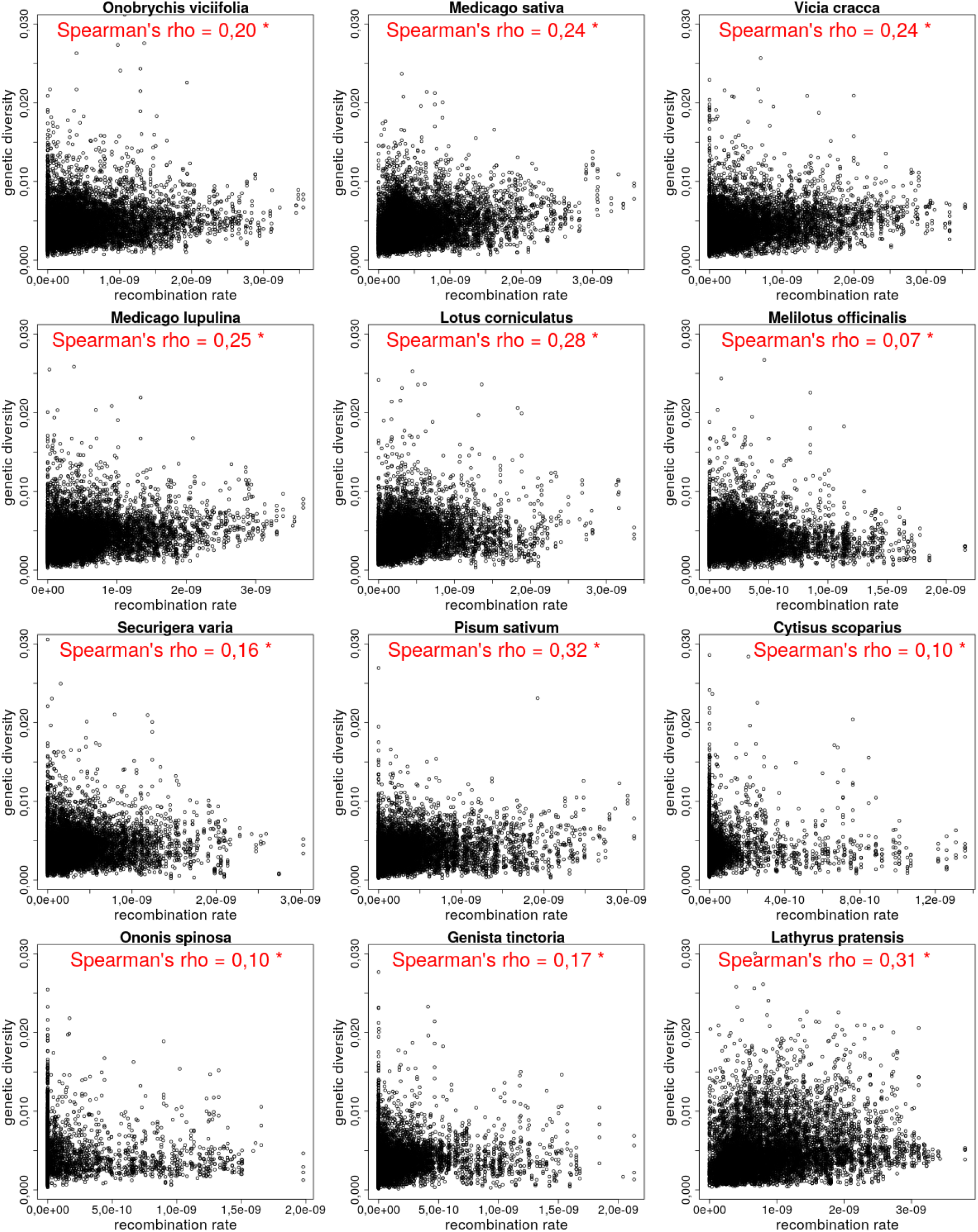
Genome-wide relationships between recombination rate and genetic diversity (*π*) across the continuum.

**Figure S10:**
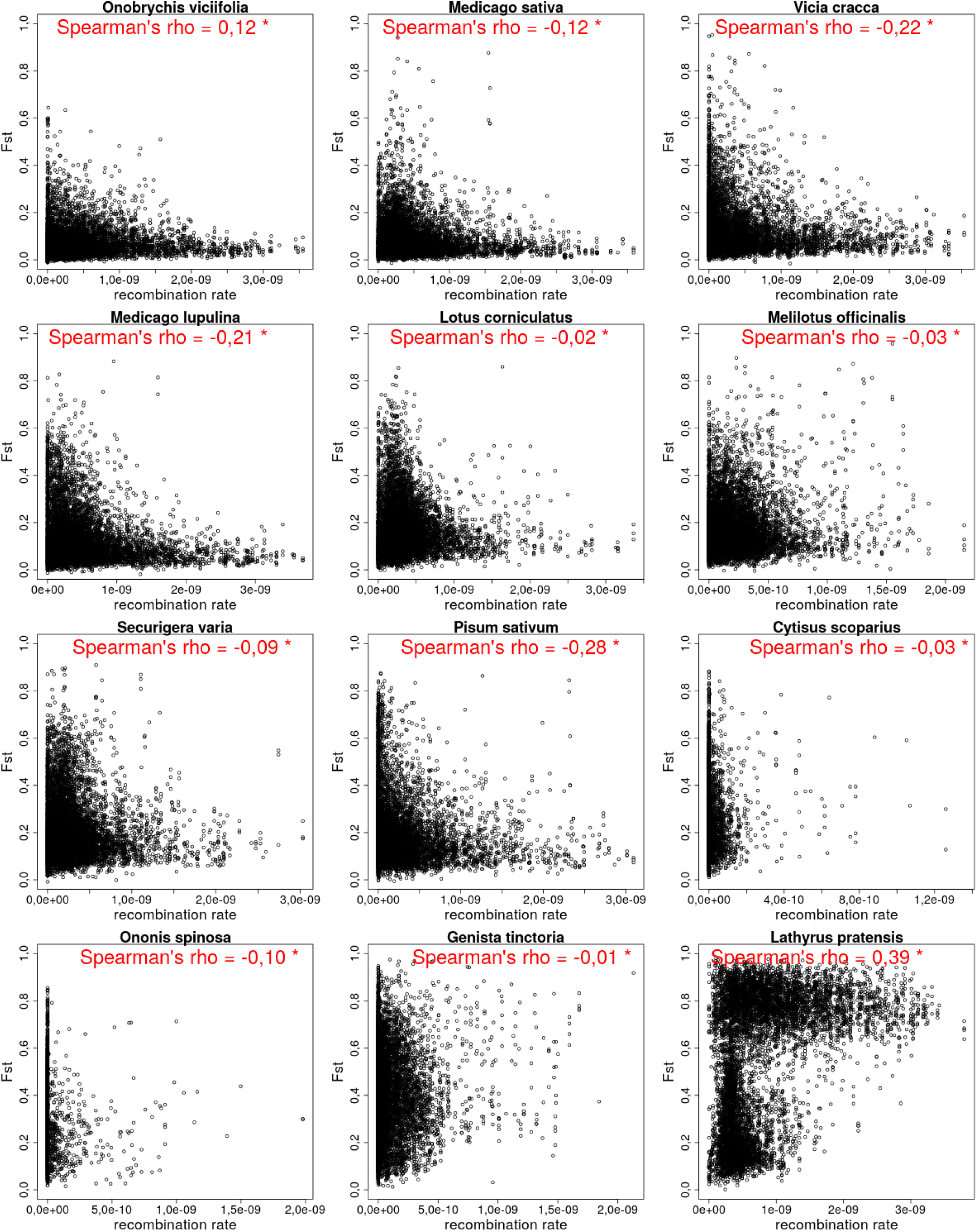
Genome-wide relationships between recombination rate and *D_XY_* across the continuum.

**Figure S11:**
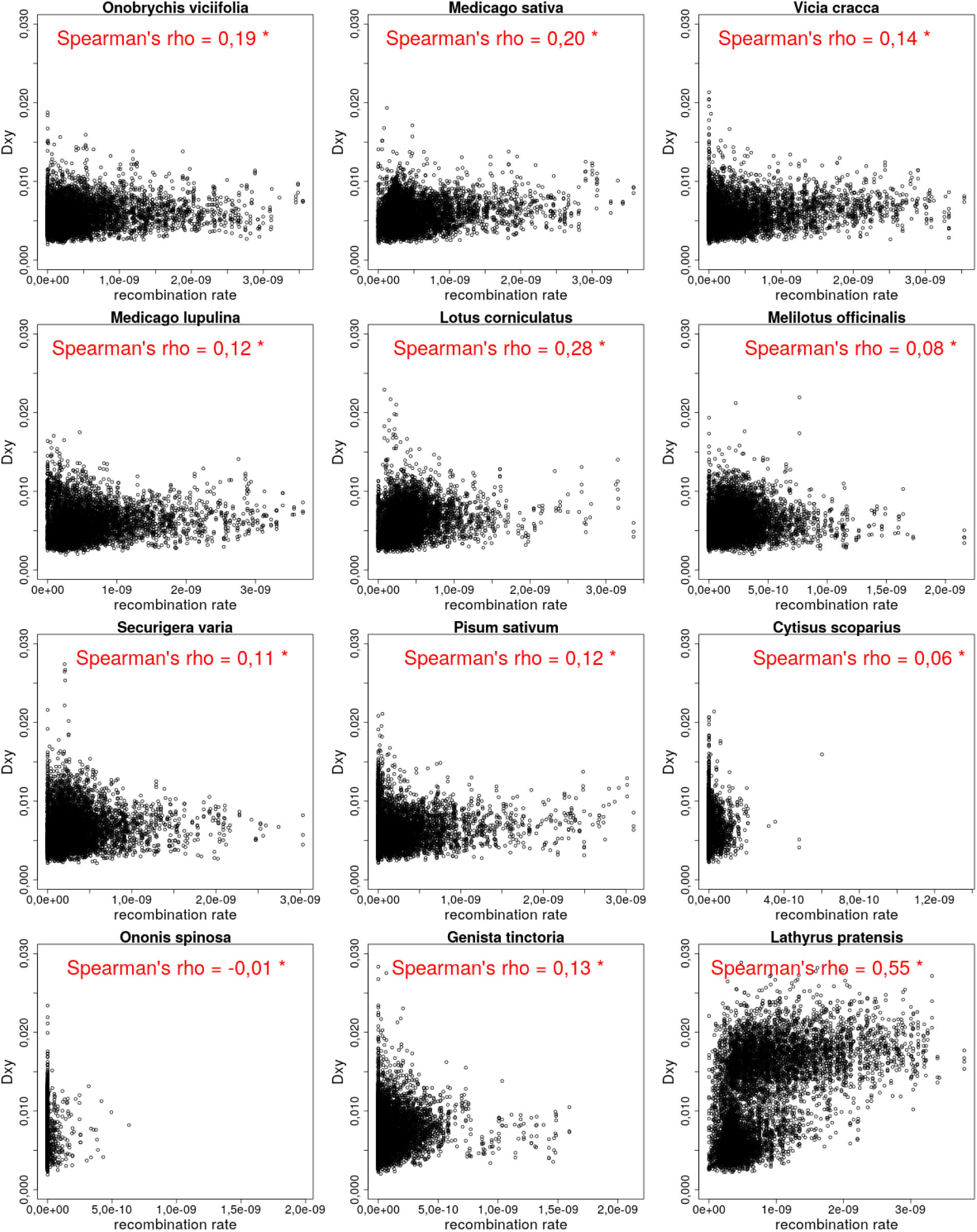
Genome-wide relationships between recombination rate and *F_ST_* across the continuum.

**Figure S12:**
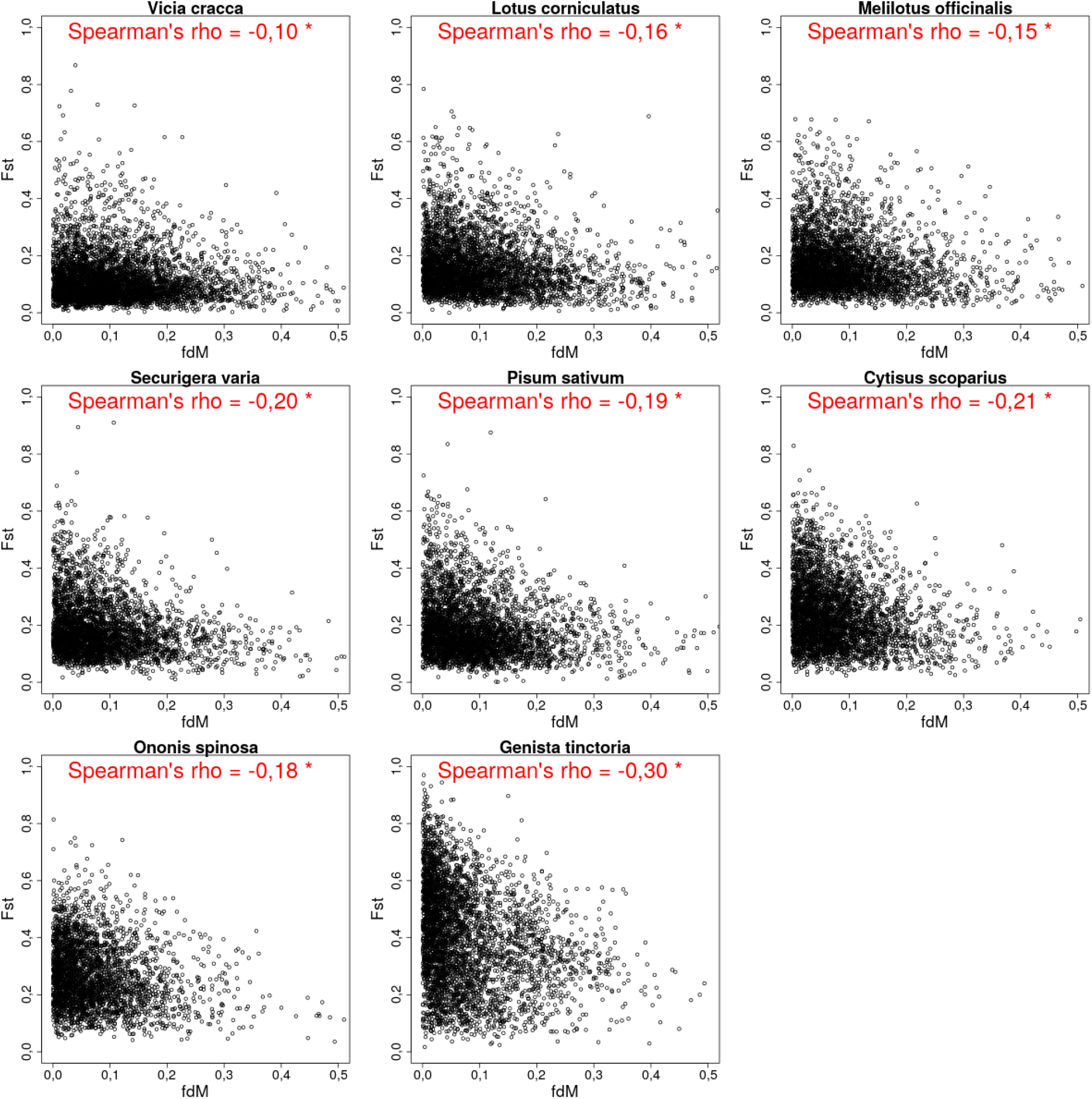
Genome-wide relationships between *F_ST_* and *fdM* across the continuum. This shows only positive fdM values.

**Figure S13:**
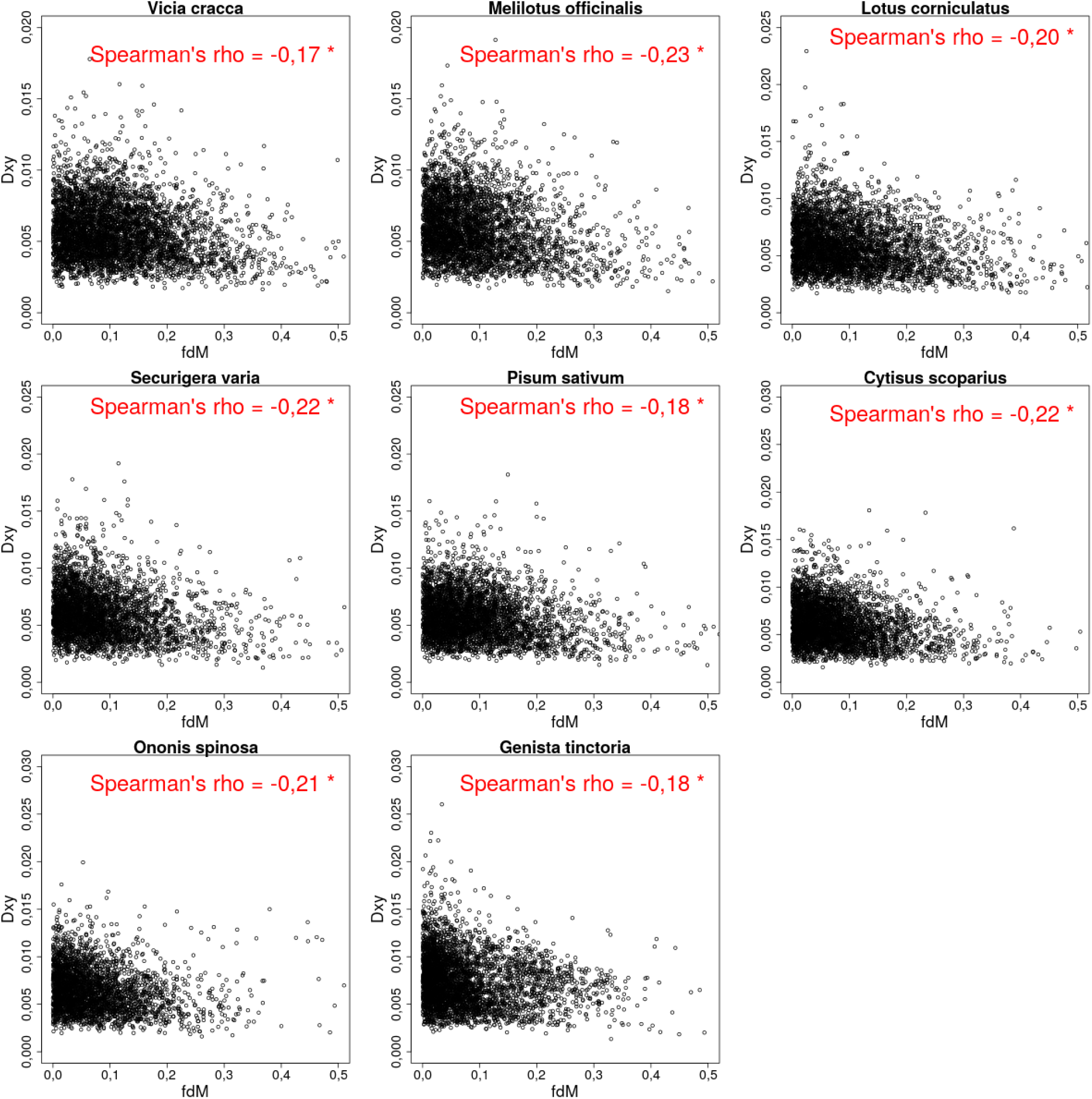
Genome-wide relationships between *D_XY_* and *fdM* across the continuum. This shows only positive fdM values.

**Figure S14:**
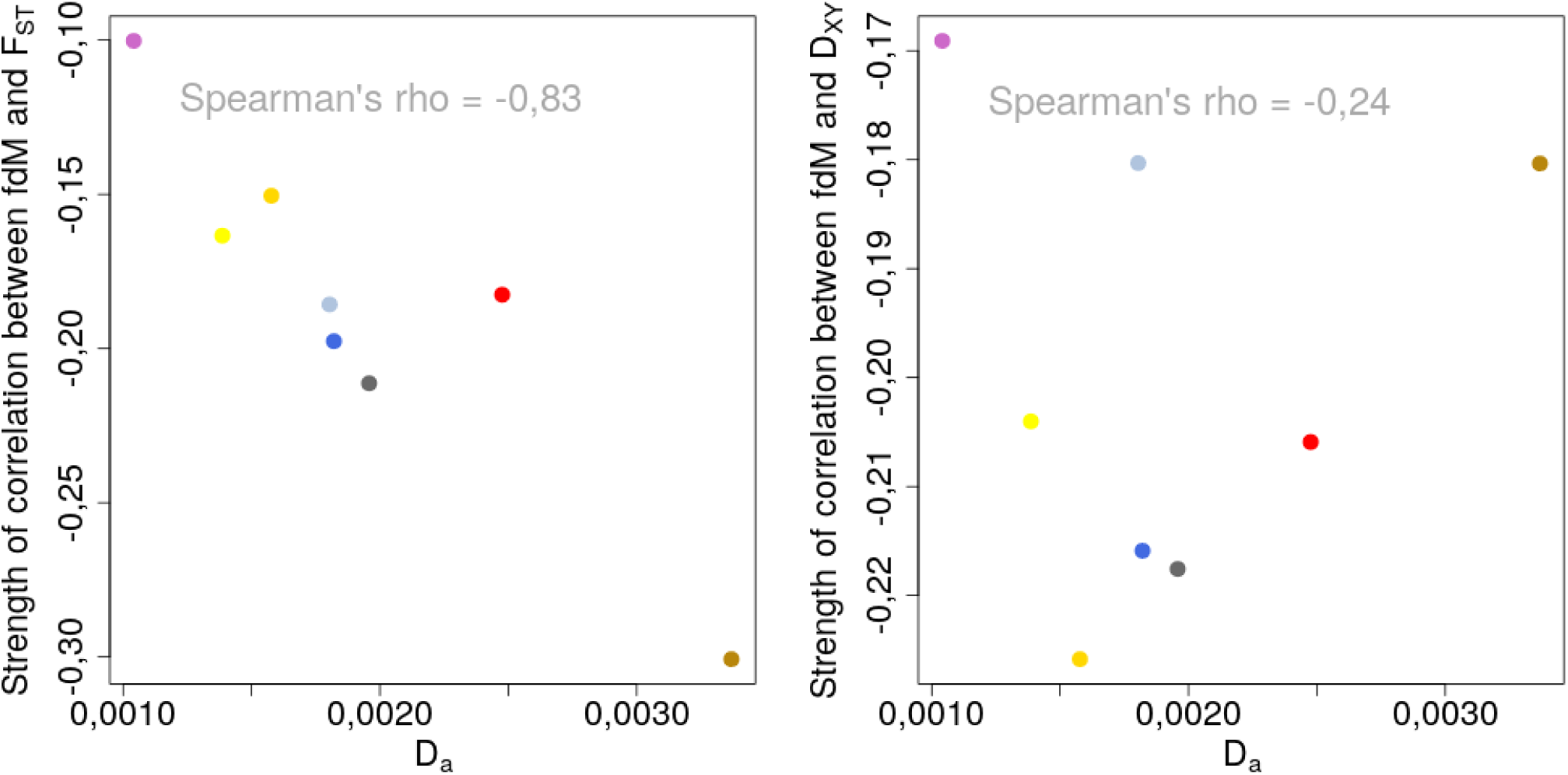
Evolution of the strengths of Spearman correlations between *fdM* and both *F_ST_* (left) and *D_XY_* (right) as divergence increases along the continuum. Colours correspond to biotypes in Figure 1A.

**Figure S15:**
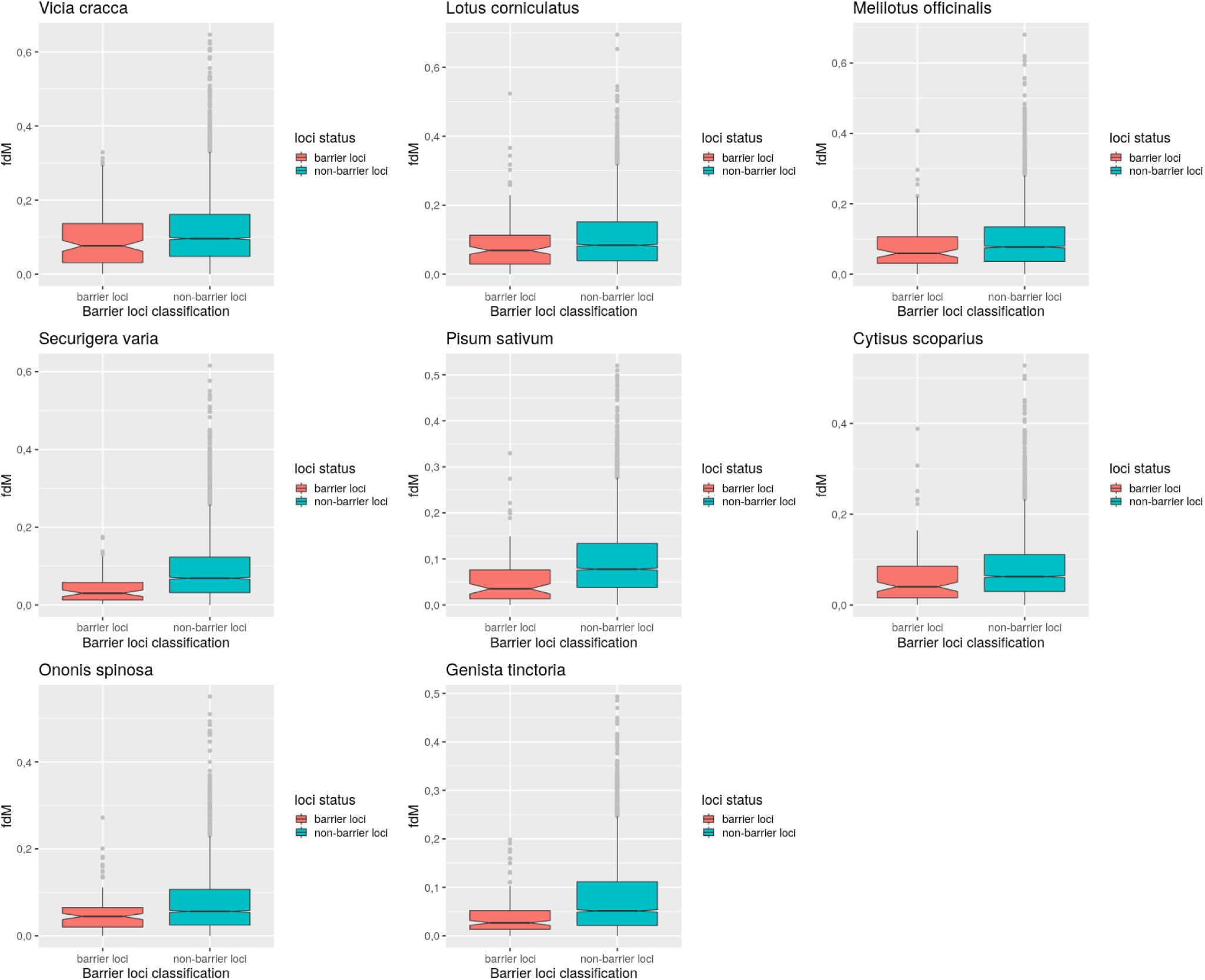
Distribution of *fdM* within and outside of inferred barrier loci. This shows only positive *fdM* values.

**Figure S16:**
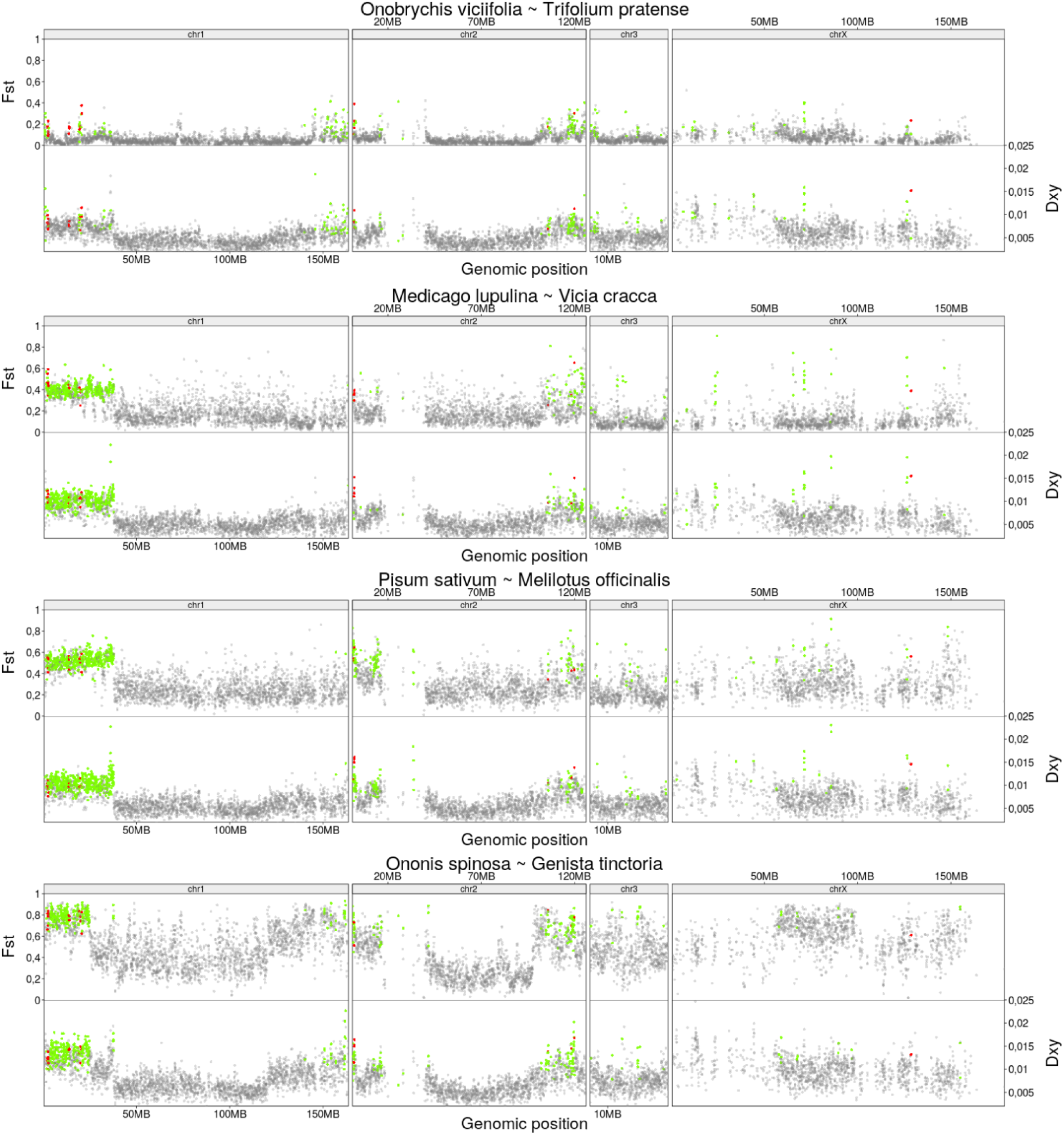
Manhattan plots of the landscapes of *F_ST_* (top), *D_XY_* (bottom) and the localisation of inferred barrier loci (green dots) and shared barrier across all pairs (red dots) in the four independent biotype pairs of increasing divergence (top to bottom).

**Figure S17:**
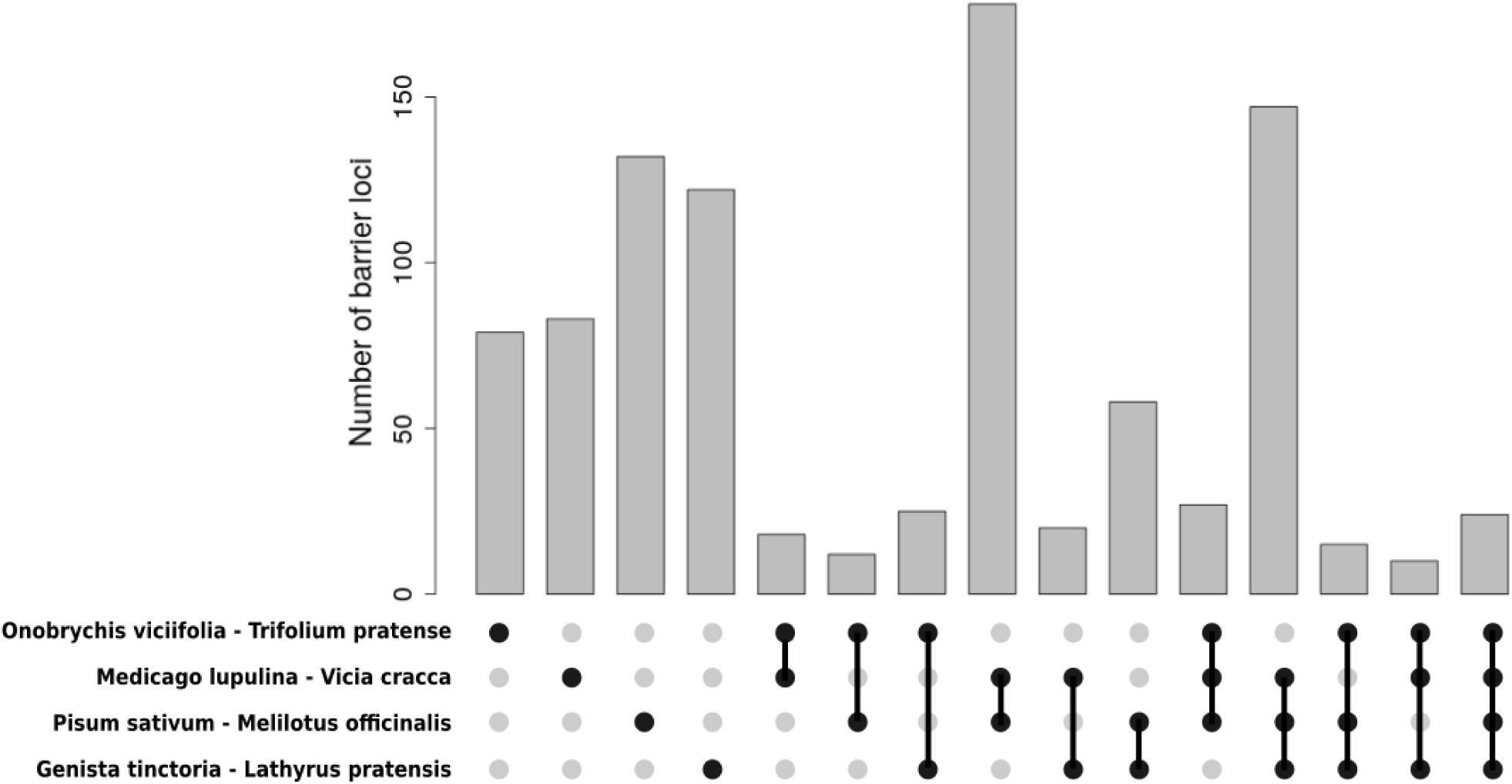
Number of unique versus shared barrier loci among the four independent pairs of biotypes. All categories are mutually exclusive.

**Figure S18:**
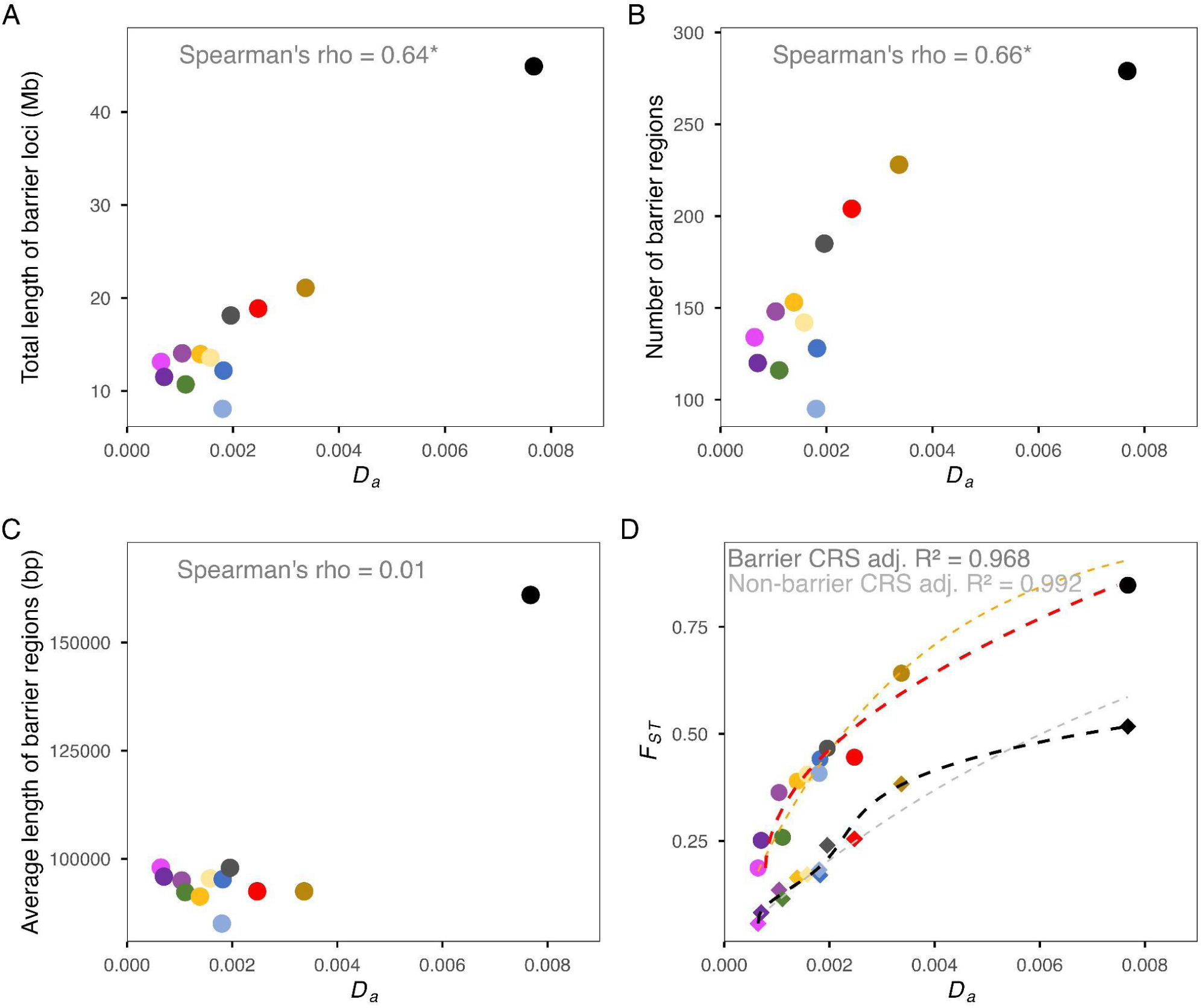
Dynamics of genomic regions acting as barriers to gene flow as divergence increases across the continuum, when candidate chromosomal rearrangements are excluded from the analyses. Coloured dots represent biotype pairs as in Figure 1. **A**: Total base pairs in barrier regions (i.e., merged adjacent barrier loci) loci as net divergence (*D_a_*) increases (in Mb), **B**: Number of (merged) barrier regions as net divergence increases, **C**: Average length of (merged) barrier regions as net divergence increases, **D**: Mean *F_ST_*within barrier windows as net divergence increase (circular points) and mean *F_ST_* in non-barrier windows (randomly chosen among all non-barrier windows to obtain the same number of non-barrier windows as the number of identified barrier windows, for each biotype pair) as divergence increases (diamond points). Red (for barrier loci) and black (for non-barrier loci) dashed lines show regression spline (CRS) fits; orange (for barrier loci) and grey (for non-barrier loci) dashed lines show asymptotic fits.

**Figure S19:**
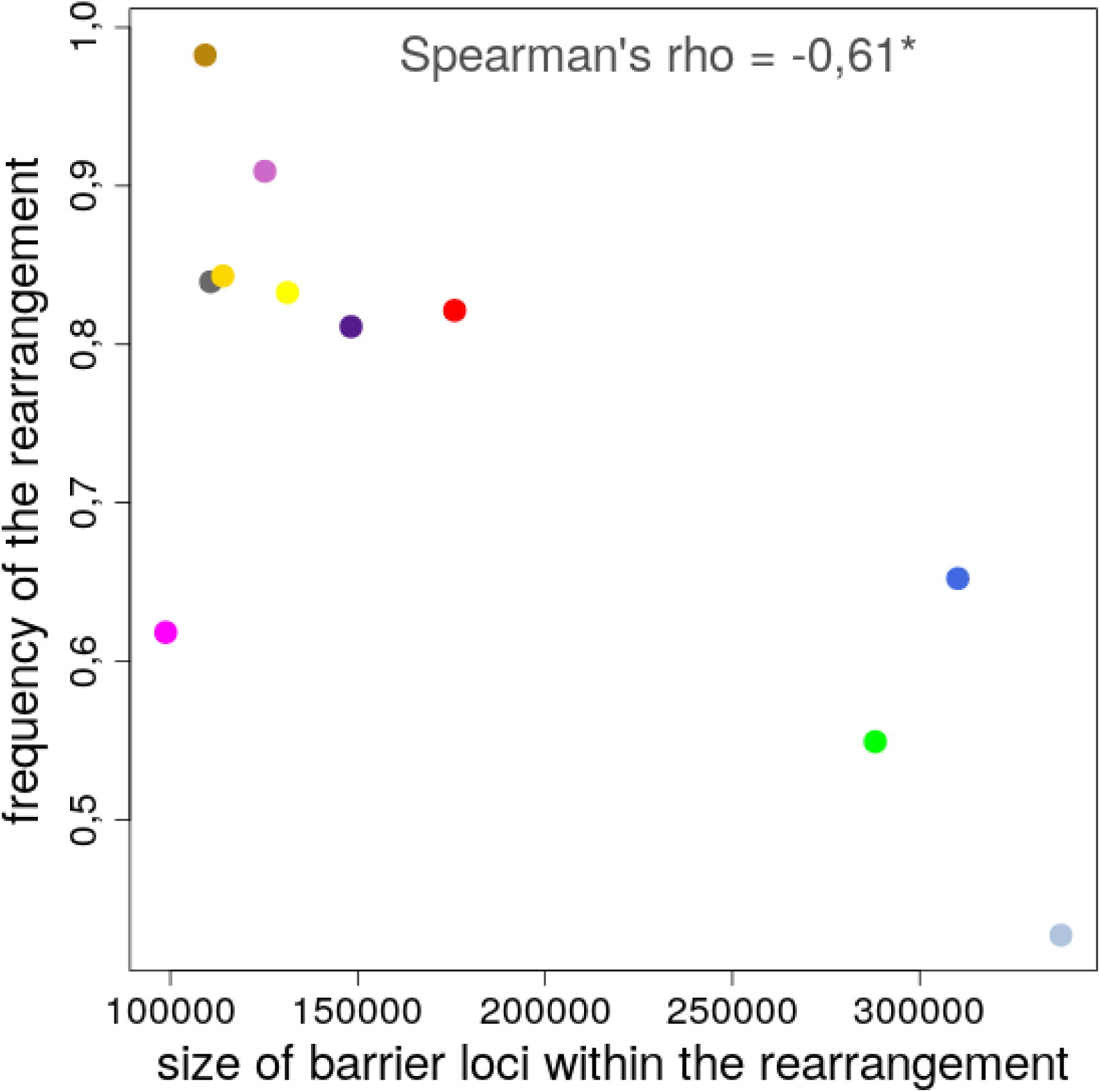
Relation between chromosome 1 structural rearrangement segregating frequency and the average size of barrier loci within it.

**Figure S20:**
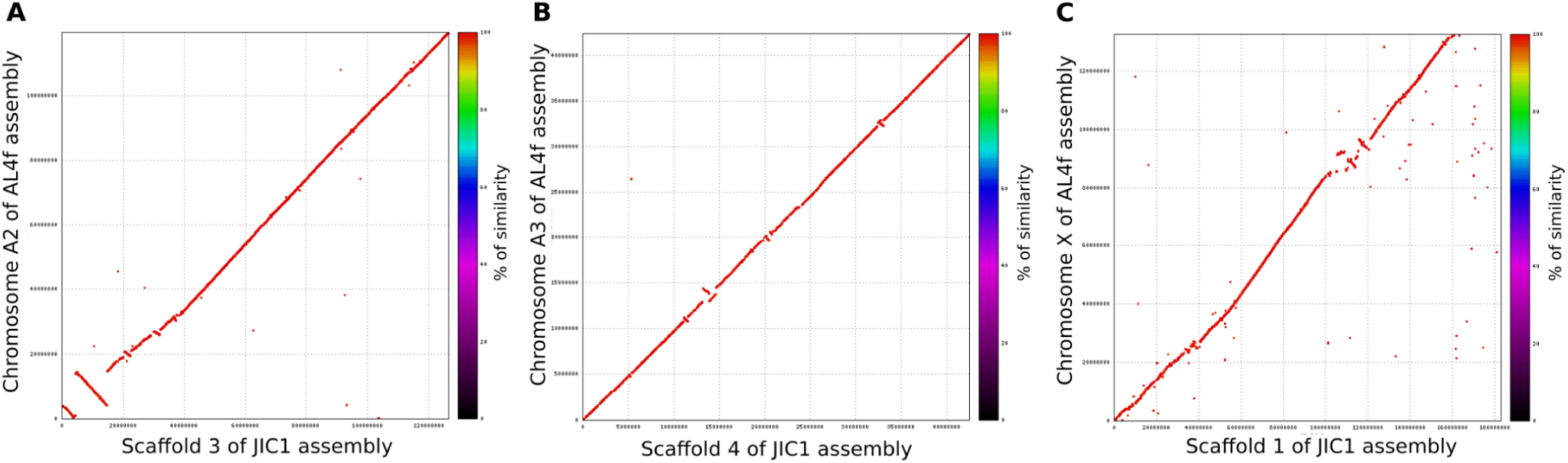
Dot plots of chromosomes A2, A3 and X from the *A. pisum* assembly based on a *Medicago sativa* individual AL4f (GCA_005508785.2) (Li et al. 2019) *versus* scaffolds 3, 4 and 1, respectively, from the *A. pisum* assembly based on a putative *Pisum sativum* individual JIC1 (Mathers et al. 2021).

**Figure S21:**
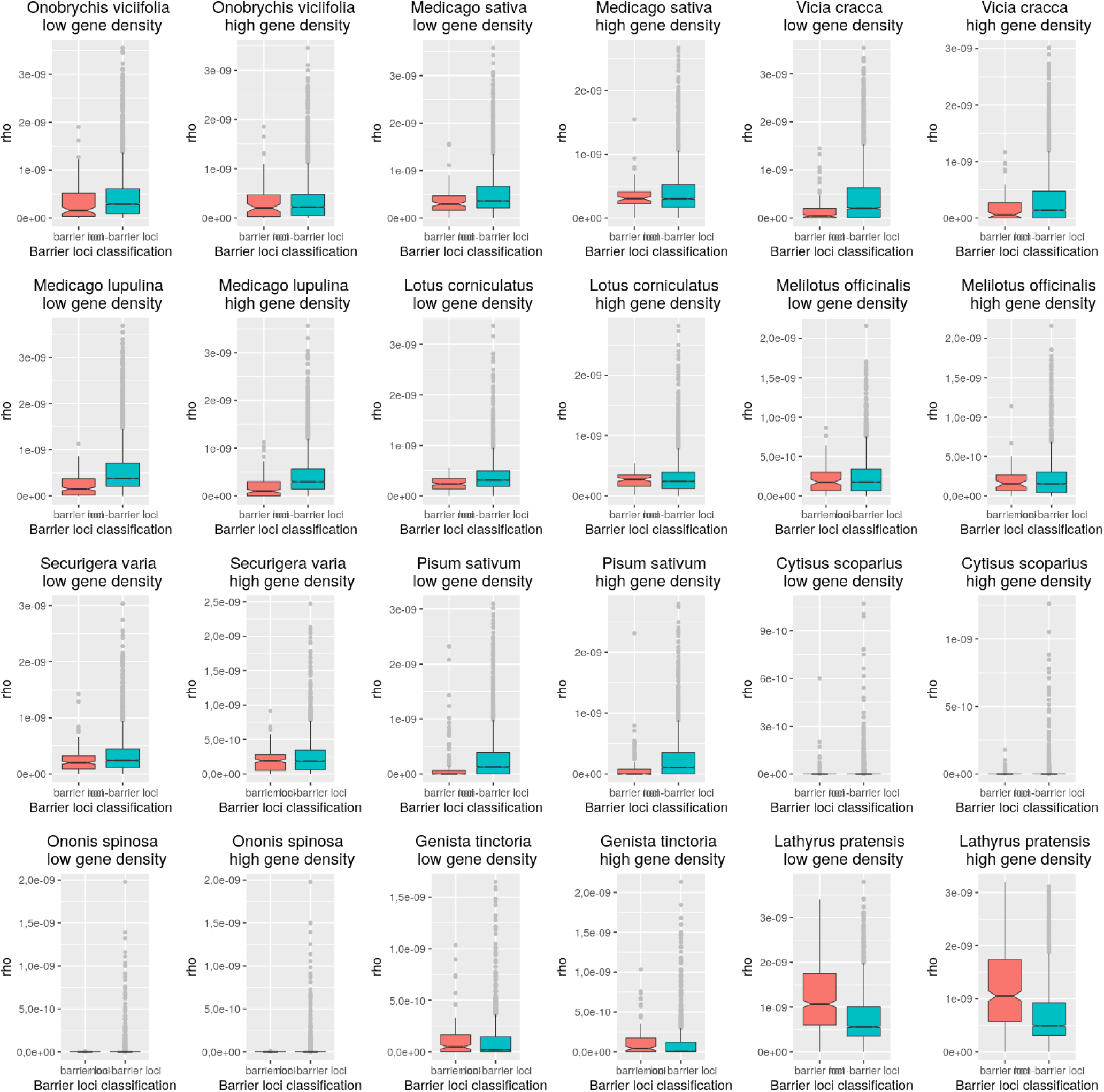
Distribution of recombination rate within and outside of inferred barrier loci in windows of low (left) *versus* high gene density (right). Significant Wilcoxon rank sum tests comparing the recombination rate within barrier loci *versus* within non-barrier loci are indicated with a star.

**Figure S22:**
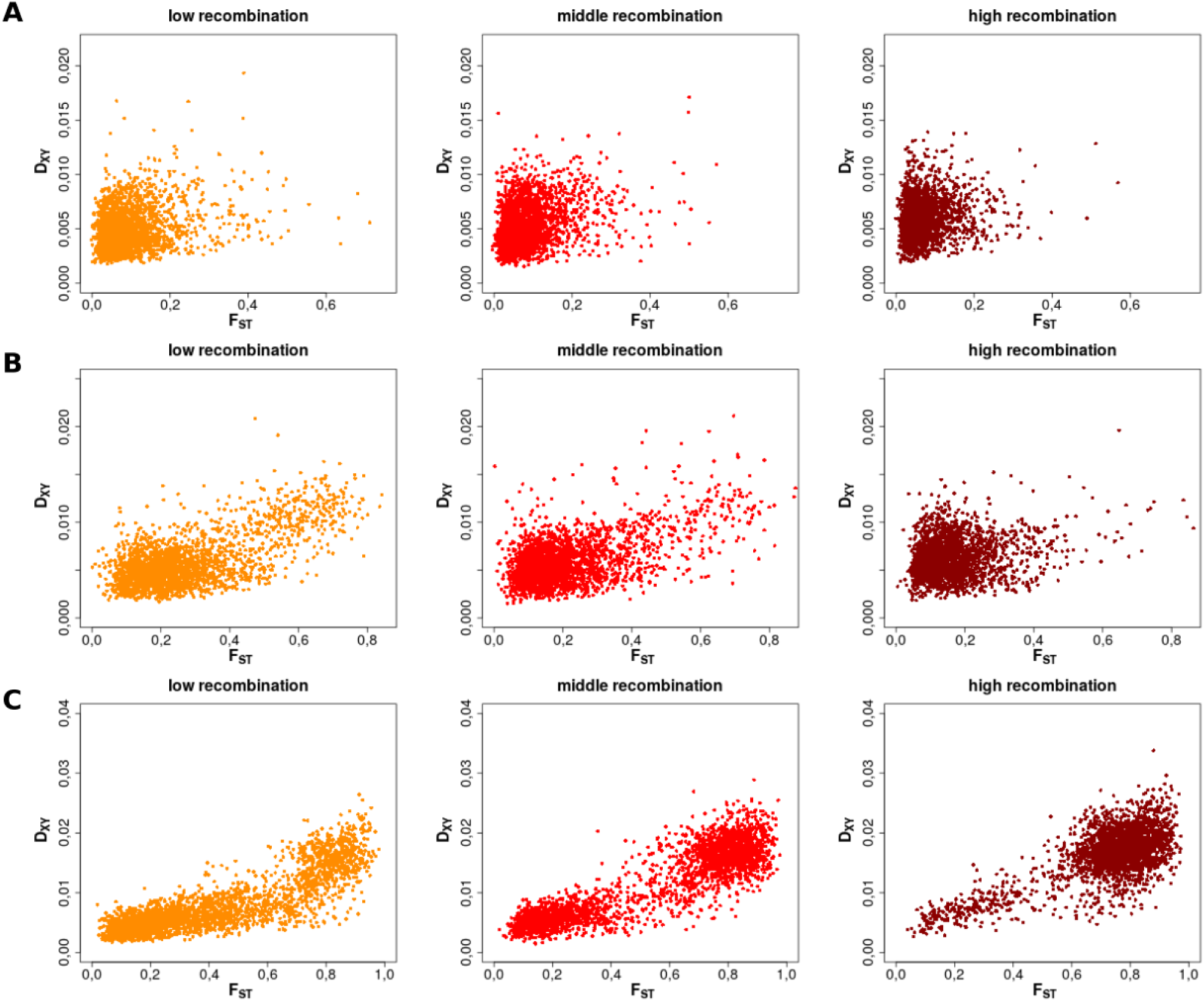
Dispersion of *F_ST_*and *D_XY_* values in bins of low, middle and high recombination rate in three biotype pairs: A: *Medicago sativa - Trifolium*, B: *Pisum sativum - Trifolium*, and C: *Lathyrus pratensis - Trifolium*.

**Figure S23:**
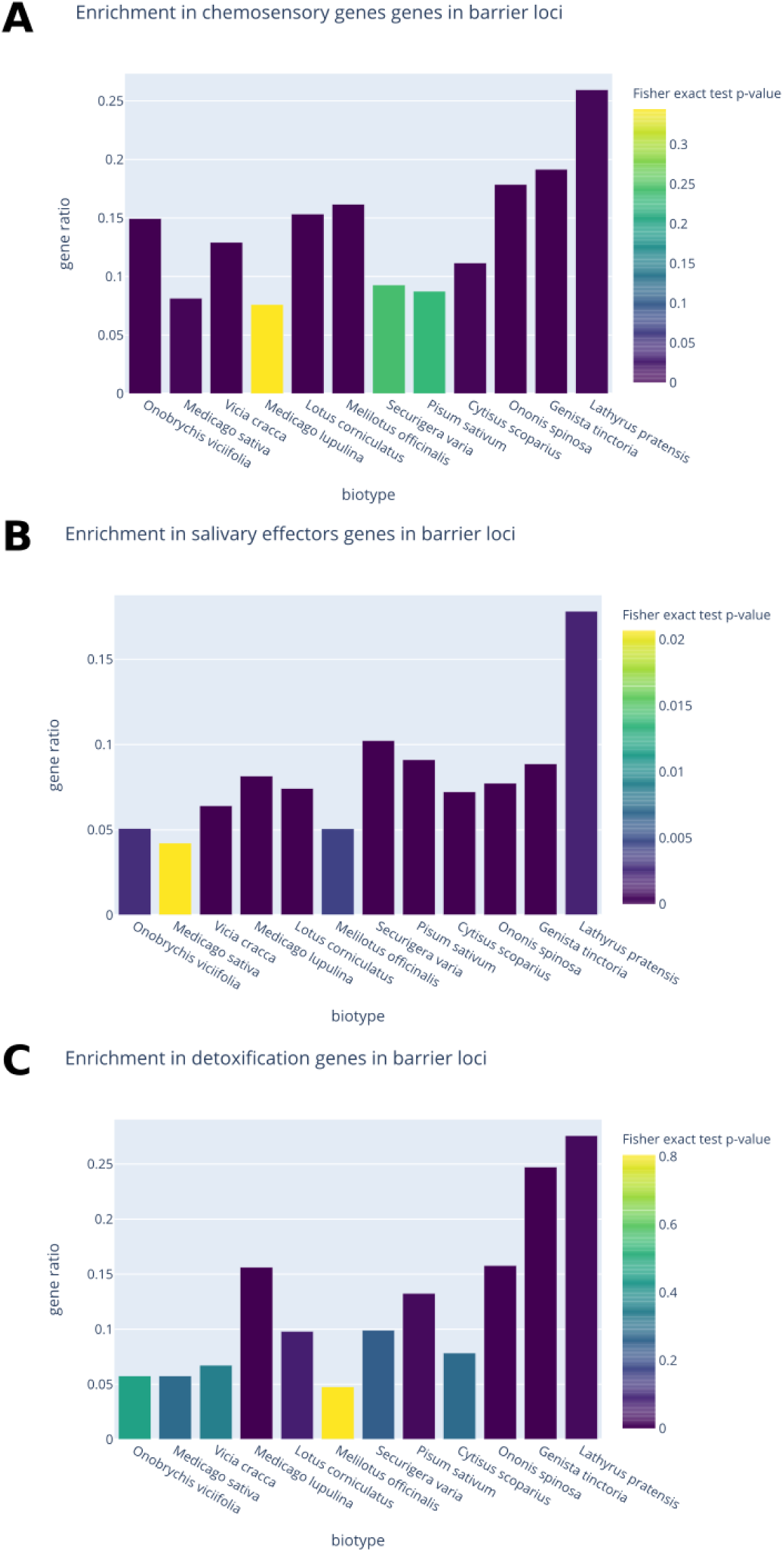
Enrichment tests of chemosensory (A), salivary effectors (B) and detoxification (C) genes within barrier loci across the continuum. Biotypes are sorted by increasing divergence with the *Trifolium* biotype.

**Figure S24:**
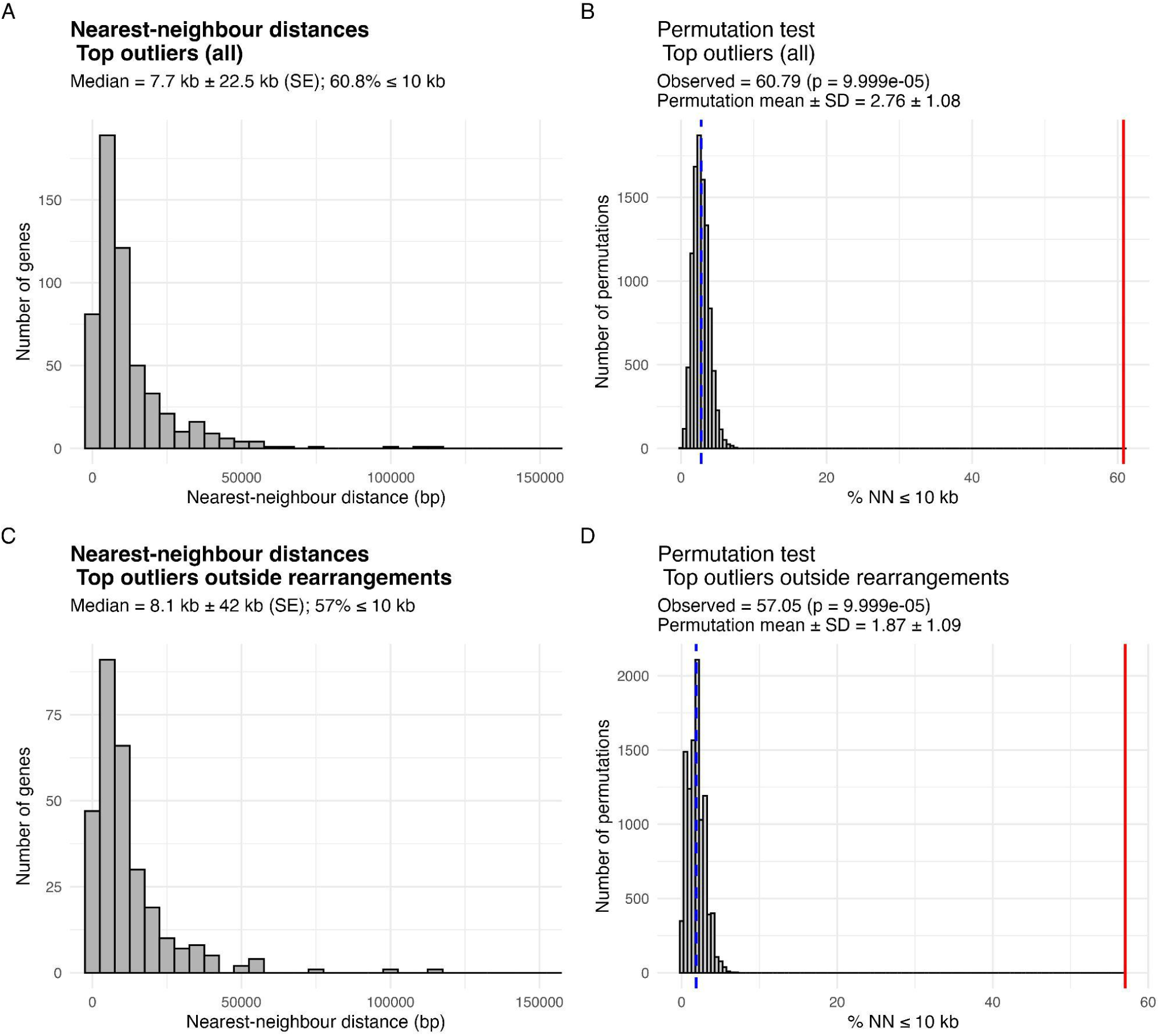
Physical clustering of top outlier genes. Panels A and C show the distribution of nearest-neighbour (NN) distances among top outlier genes, with grey bars representing counts of genes in each 5 kb bin, and red vertical lines indicating the observed NN statistic. Panels B and D show the distribution of permutation statistics across 10,000 randomisations, with grey bars representing the random expectation and the red vertical line showing the observed statistic. Panels A–B include all top outliers, while C–D include only top outliers outside candidate chromosomal rearrangements. The 10 kb threshold indicates the a priori definition of “close” genes.

**Figure S25:**
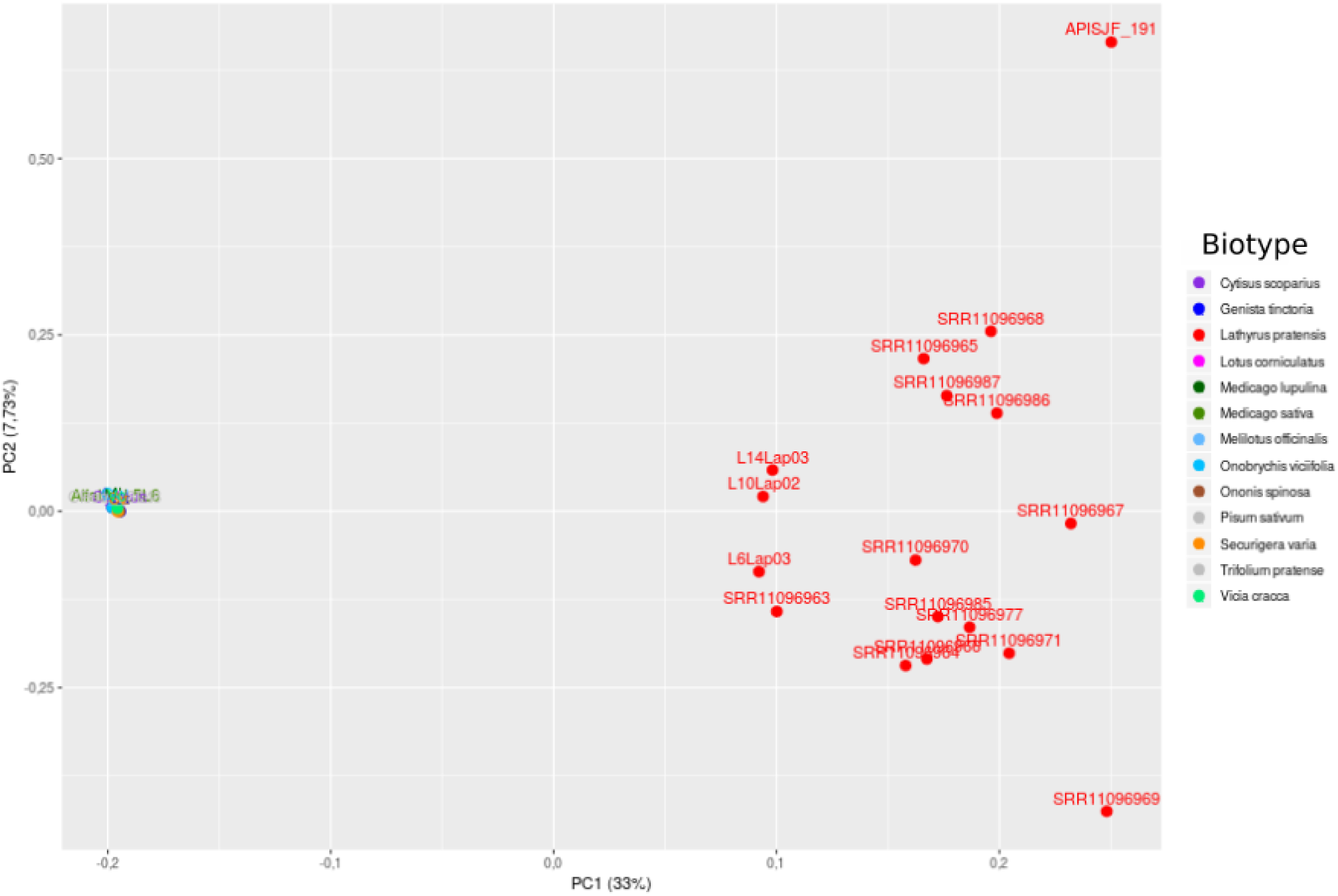
PCA of individual *Lathyrus pratensis* samples subsequently pooled *versus* pooled samples of the rest of the biotypes.

**Figure S26:**
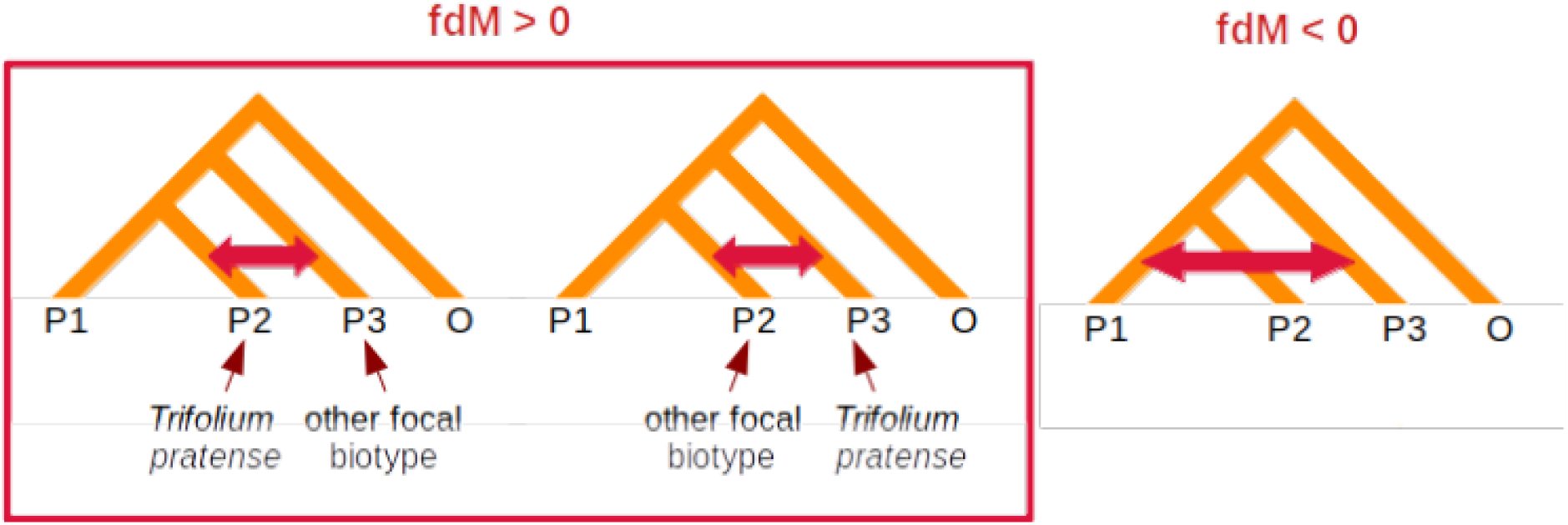
Schematic of the reasoning of the computation of the *fdM* statistic in the context of the continuum.

**Figure S27:**
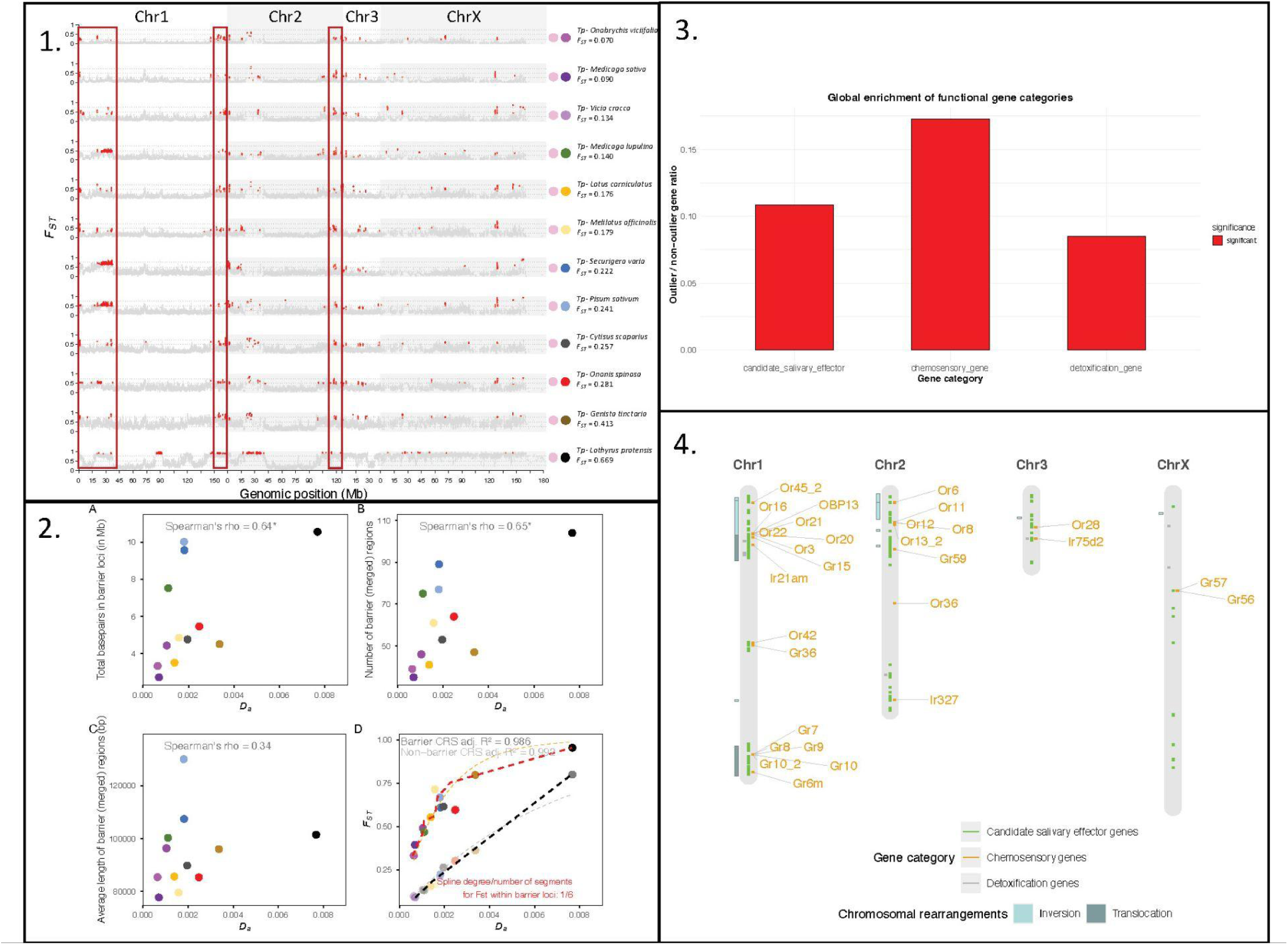
Reproduction of the main results when applying FDR correction on *p*-values for *F_ST_* and *D_XY_* before identifying barrier loci. (1) Equivalent to Figure 3; (2) Equivalent to Figure 4; (3) Functional enrichment results for the three candidate gene categories; (4) Equivalent to Figure 6.

